# Early lock-in of structured and specialised information flows during neural development

**DOI:** 10.1101/2021.06.29.450432

**Authors:** David P. Shorten, Viola Priesemann, Michael Wibral, Joseph T. Lizier

## Abstract

The brains of many organisms are capable of complicated distributed computation underpinned by a highly advanced information processing capacity. Although substantial progress has been made towards characterising the information flow component of this capacity in mature brains, there is a distinct lack of work characterising its emergence during neural development. This lack of progress has been largely driven by the lack of effective estimators of information processing operations for the spiking data available for developing neural networks. Here, we leverage recent advances in this estimation task in order to quantify the changes in information flow during development. We do so by studying the changes in the intrinsic dynamics of the spontaneous activity of developing dissociated neural cell cultures. We find that the quantity of information flowing across these networks undergoes a dramatic increase across development. Moreover, the spatial structure of these flows is locked-in during early development, after which there is a substantial temporal correlation in the information flows across recording days. We analyse the flow of information during the crucial periods of population bursts. We find that, during these bursts, nodes undertake specialised computational roles as either transmitters, mediators or receivers of information, with these roles tending to align with their spike ordering — either early, mid or late in the bursts. Further, we find that the specialised computational roles occupied by nodes during bursts tend to be locked-in early. Finally, we briefly compare these results to information flows in a model network developing according to an STDP learning rule from a state of independent firing to synchronous bursting. The phenomena of large increases in information flow, early lock-in of information flow spatial structure and computational roles based on burst position were also observed in this model, hinting at the broader generality of these phenomena.

**AUTHOR SUMMARY:** This paper studies the development of computation in biological systems by analysing changes in the flow of information in developing neural cell cultures. Although there have been a number of previous studies of information flows in neural cell cultures, this work represents the first study which compares information flows in the intrinsic dynamics across development time. Moreover, we make use of a recently proposed continuous-time transfer entropy estimator for spike trains, which, in comparison to the discrete-time estimator used previously, is able to capture important effects occurring on both small and large timescales simultaneously. We find that information flows begin to emerge after 5-10 days of activity, and crucially, the spatial structure of information flows remains significantly temporally correlated over the first month of recording. Furthermore, the magnitude of information flows across the culture are strongly related to burst position, and the roles of regions as information flow sources, sinks and mediators are found to remain consistent across development. Finally, we confirm that these early lock-ins also occur in a simple model network developing under an STDP update rule, suggesting a plausible mechanism undergirding this phenomenon.

## I. INTRODUCTION

Throughout development, how do brains gain the ability to perform advanced computation? Given that the distributed computations carried out by brains require an intrinsic information processing capacity, it is of utmost importance to decipher the nature of the emergence of this capacity during development.

For brains to engage in the computations required for specific tasks, they require a general-purpose computational *capacity*. This capacity is often studied within the framework of information dynamics, where it is decomposed into the atomic operations of information storage, transfer and modification [1, 2]. We are particularly interested in the information flow component, which is measured using the Transfer Entropy (TE) [3, 4]. There exists a substantial body of work examining the structure and role of computational capacity in terms of these operations in mature brains. This includes: the complex, dynamic, structure of information transfer revealed by calcium imaging [5], fMRI [6, 7], MEG [8] and EEG [9–12], and the role of information storage in representing visual stimuli [13], among others. Given the established role of information flows in enabling the computations carried out by mature brains, we aim to study how they self-organise during neural development. There are a number of requirements for such a study. Firstly, it needs to be performed at a fine spatial scale (close to the order of individual neurons), to capture the details of development. It also needs to be conducted longitudinally in order to track changes over developmental timescales. Finally, the estimation of the information flow as measured by TE needs to be performed with a technique which is both accurate and able to capture the subtleties of computations performed on both fine and large time scales simultaneously.

Considering the first requirement of fine spatial scale, cell cultures plated over Multi-Electrode Arrays (MEAs) allow us to record from individual neurons in a single network, providing us with this fine spatial resolution. There have been a number of previous studies examining information flows in neural cell cultures, e.g.: [14–20]. This work has focussed on the functional networks implied by the estimated TE values between pairs of nodes which has revealed interesting features of the information flow structure. See Sec. IV D 1 for a more detailed description of this previous work.

However, moving to our second requirement of a longitudinal study, these studies have almost exclusively examined only single points in neural development, since nearly all of them examined recordings from slice cultures of mature networks. By contrast, we aim to study the information flows longitudinally, by estimating them at different stages in development. Using recordings from developing cultures of dissociated neurons [21] makes this possible.

In terms of our third requirement of accurate and high-fidelity estimation of TE, we note that all previous studies of information flows in neural cell cultures made use of the traditional discrete-time estimator of TE. As recently demonstrated [22], the use of this estimator is problematic, as it can only capture effects occurring on a single time-scale. In contrast, a novel continuous-time TE estimator [22] captures effects on multiple scales, avoiding time-binning, is data efficient and consistent. See Sec. IV D for a more detailed discussion of the differences between the continuous-time and discrete-time estimators.

In this paper, we thus examine the development of neural information flows for the first time, addressing the above requirements by applying the continuous-time TE estimator to recordings of developing dissociated cultures. We find that the amount of information flowing over these cultures undergoes a dramatic increase throughout development and that the patterns of these flows are established early. During bursting periods we find that nodes engage in specialised computational roles as either transmitters, receivers or mediators of information flow. Moreover, these roles correspond with the node’s position in the burst propagation, with middle bursters tending to be information mediators. This provides positive evidence for the pre-existing conjecture that nodes in the middle of the burst propagation play the vital computational role of “brokers of neuronal communication” [23]. Intriguingly, the designation of computational roles (transmitter, receiver or mediator) appears to be determined early in development. Finally, in order to investigate the generality of these phenomena, as well as a putative mechanism for their emergence, we study the dynamics of information flow in a model network developing according to an STDP update rule. We find that the above-mentioned phenomena are present in this model system, hinting at the broader generality of such patterns of information flow in neural development.

## II. RESULTS

Data from overnight recordings of developing cultures of dissociated cortical rat neurons at various stages of development (designated by days in vitro, DIV) was analysed. These recordings are part of an open, freely available, dataset [21, 24]. See methods (Sec. IV A) for a summary of the setup that produced the recordings. We selected four cultures from the dataset to study, which we refer to by the same naming convention used in the open dataset: 1-1, 1-3, 2-2 and 2-5. Each culture has overnight recordings at four different time points, apart from 1-1, which was only recorded thrice. The days on which these recordings took place vary between the 4th and 33rd DIV. By contrasting the TE values estimated at these different recording days, we are able to obtain snapshots of the emergence of these culture’s computational capacity.

The TE between all pairs of electrodes was estimated using a recently introduced continuous-time estimator [22] (see Sec. IV D). This produces a directed functional network at each recording day, and we aim to analyse how the connections in this network change over development time. Spike sorting was not performed, because we would not be able to match the resulting neural units across different recordings, and could not then fulfil our aim of contrasting the information flow between specific source-target pairs at different recording days. As such, the activity on each node in the directed functional networks we study is multi-unit activity (MUA) [23] formed of the spikes from all neurons detected by a given electrode, with connections representing information flows in the MUA. For more detail on data pre-processing as well as the parameters used with the estimator, see Methods (Sec. IV).

### A. The dramatic increase in the flow of information during development

We first investigate how the amount of information flowing between the nodes changes over the lifespan of the cultures. Table I shows the mean TE between all source-target pairs. We observe that this mean value increases monotonically with the number of DIV, with only a single exception (a slight drop in the mean TE between days 15 and 21 of culture 2-2). Otherwise, the magnitude of the increase in the mean TE is substantial. Among the first recordings for each culture, both recordings on the 4th DIV had a mean estimated TE of 0 nats.s^−1^ (with no statistically significant transfer entropies measured as per Sec. II B), the single recording on the 5th DIV had a mean of 0.013 nats.s^−1^ and the single recording on the 9th DIV had a mean of 0.0084 nats.s^−1^. By contrast, all recordings beyond 20 DIV had a mean TE greater than 0.049 nats.s^−1^ and all recordings beyond 28 DIV had a mean TE greater than 0.15 nats.s^−1^.

**TABLE I:** Mean TE in nats per second between every source-target pair for each recording studied.

|  |  |  |  |  |
| --- | --- | --- | --- | --- |
| **Culture 1-1** | day 4 | day 14 | day 20 |  |
|  | 0 | 0.018 | 0.058 |  |
| **Culture 1-3** | day 5 | day 10 | day 16 | day 24 |
|  | 0.013 | 0.016 | 0.020 | 0.068 |
| **Culture 2-2** | day 9 | day 15 | day 21 | day 33 |
|  | 0.0084 | 0.066 | 0.049 | 0.15 |
| **Culture 2-5** | day 4 | day 10 | day 22 | day 28 |
|  | 0 | 0.040 | 0.10 | 0.19 |

Fig. 1a shows scatter plots of the TE values in each recording laid over box-and-whisker plots. The large increase over time in the amount of information flowing over the networks is clearly visible in these plots. However, it is interesting to note that certain source-target pairs do have large information flows between them on early recording days even whilst the average remains very low.

**FIG. 1:**
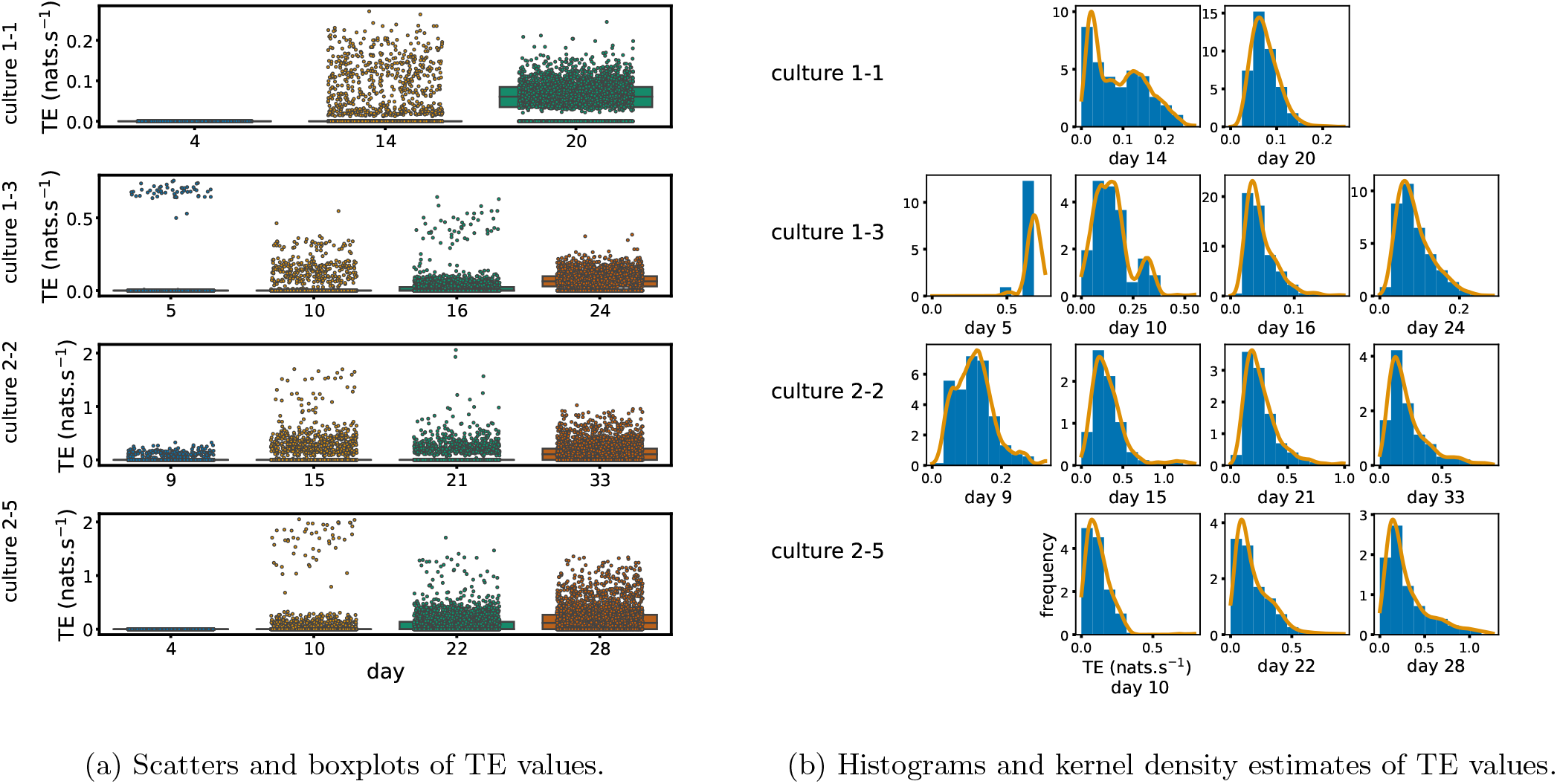
Plots of the distributions of estimated TE values in the recordings analysed in this study. (a) Scatters of the TE values are overlaid on box plots. The box plots show the quartiles and the median (values greater than 10 standard deviations from the mean have been removed from both the box and scatter plots as outliers). (b) Density estimates of the nonzero (statistically significant) TE distribution on top of a histogram. The densities are estimated using a Gaussian kernel. The histogram bin width and kernel histogram are both 10% of the data range.

Fig. 1b shows histograms of the TE values estimated in each recording along with probability densities estimated using a Gaussian kernel. The distributions only include the nonzero (statistically significant) estimated TE values. These distributions do, qualitatively, appear to be log-normal, in particular for later DIV. Moreover, previous studies have placed an emphasis on the observation of log-normal distributions of TE values in *in vitro* cultures of neurons [14, 15]. As such, we qantitatively analysed the distribution of the nonzero (statistically significant) estimated TE values in each individual recording. However, contrary to expectations, we found that these values were not well described by a log-normal distribution. See Appendix A for further details and discussion.

### B. The emergence of functional information flow networks

By considering each electrode as a node in a network, we can construct functional networks of information flow by assigning a directed edge between each source-target pair of electrodes with a statistically significant information flow. This results in weighted networks, the weight being provided by the TE value. Diagrams of these networks are show in Fig. 2.

**FIG. 2:**
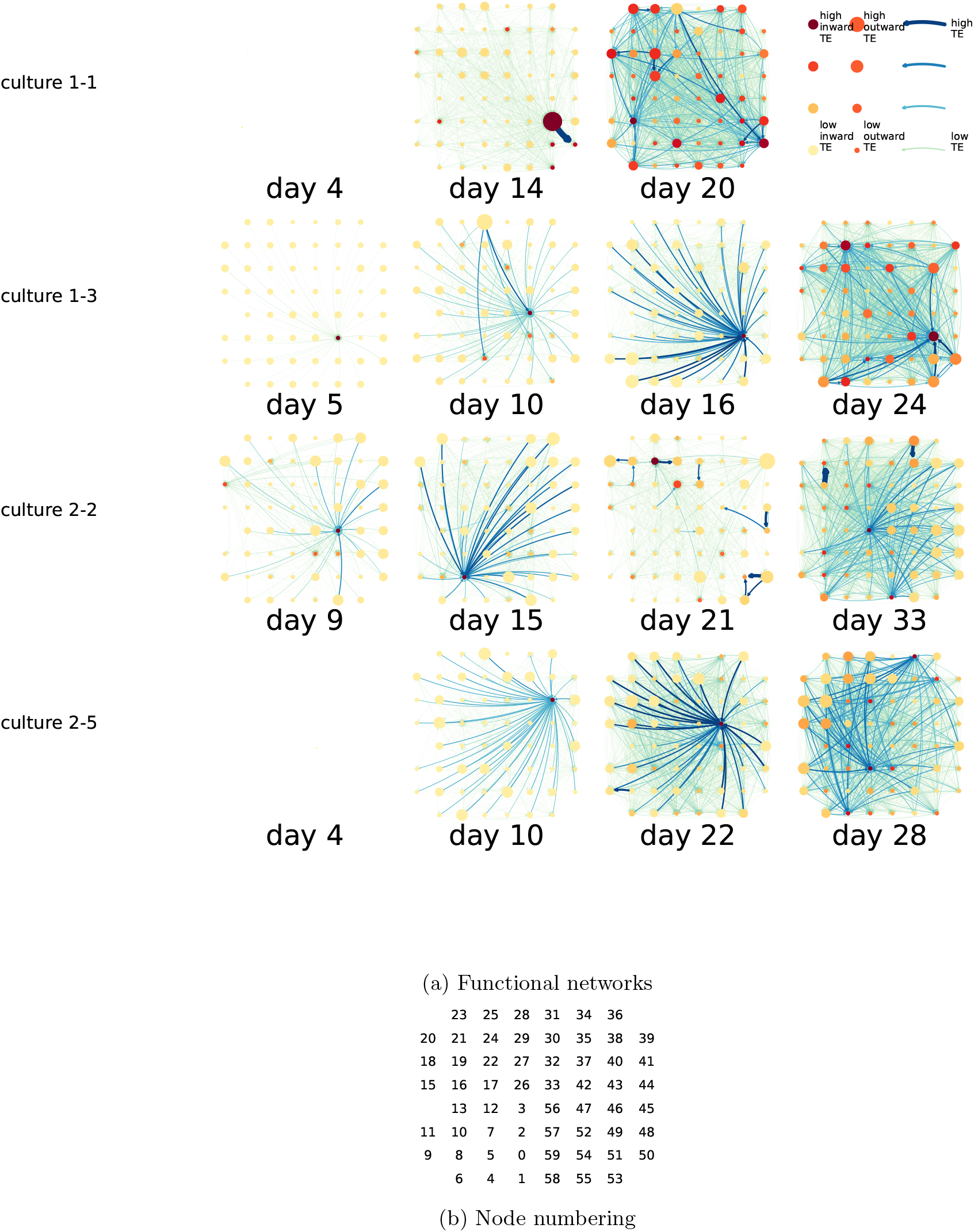
(a) The functional networks implied by the estimated TE values. Each node represents an electrode in the original experimental setup. The nodes are spatially laid out according to their position in the recording array. An edge is present between nodes if there is a statistically significant information flow between them. The edge weight and colour is indicative of the amount of information flowing between electrodes (see the legend). The scaling of this weight and colour is done relative to the mean and variance of the information flow in each recording separately. The size and colour of the nodes is assigned relative to the total outgoing and incoming information flow on the node, respectively. As with the edge colour and size, this is done relative to the distribution of these values in each recording separately. (b) The spatial layout of the nodes. The numbering is identical to that used in the documentation of the open dataset studied in this work [21, 24]

We are able to notice a number of interesting spatio-temporal patterns in these diagrams. Firstly, the density (number of edges) of the networks increases over time. This is quantified in Table II, which shows the number of source-target pairs of electrodes for which a statistically significant non-zero TE value was estimated. In all cultures studied in this work, the number of such pairs (and, therefore, the network density), increased by orders of magnitude over the life of the culture. For instance, in both cultures 1-1 and 2-5, no statistically significant TE values are estimated on the first recording day. However, around 2000 source-target pairs have significant TE values between them on the final day of recording for each culture. We are, therefore, observing the networks moving from a state where no nodes are exchanging information, to one in which information is being transferred between a substantial proportion of the pairs of nodes (≈ 58% density of possible directed connections in the network). Put another way, the functional networks are emerging from an unconnected state to a highly connected state containing the information flow structure that underpins the computational capacity of the network. This helps to explain the overall increase in information flow across the network reported in Sec. II A.

**TABLE II:** The number of source-target pairs of electrodes with a statistically significant TE value between them for each recording studied. This corresponds to the number of possible edges in the functional networks shown in Fig. 2. As the electrode arrays used to record the data had 59 electrodes, the total number of unique ordered pairs of electrodes (and, therefore, the number of possible edges) is 3422.

|  |  |  |  |  |
| --- | --- | --- | --- | --- |
| **Culture 1-1** | day 4 | day 14 | day 20 |  |
|  | 0 | 614 | 2392 |  |
| **Culture 1-3** | day 5 | day 10 | day 16 | day 24 |
|  | 55 | 311 | 823 | 2367 |
| **Culture 2-2** | day 9 | day 15 | day 21 | day 33 |
|  | 211 | 558 | 538 | 1984 |
| **Culture 2-5** | day 4 | day 10 | day 22 | day 28 |
|  | 0 | 351 | 1481 | 1942 |

We observe that the information flow (both incoming and outgoing) is spread somewhat evenly over the networks - in the sense that in the later, highly-connected, recordings there are very few areas with neither incoming nor outgoing flow. A number of clear hubs do stand out against this strong background information flow however. The strongest such hubs (with many high-TE edges) are all information sinks: they have low outgoing information flow, but receive high flow from a number of other nodes.

One can observe many instances in these diagrams where nodes have either very high incoming flow and very low outgoing flow, or very low incoming flow and very high outgoing flow. That is, they are taking on the roles of source (information-transmitting) hubs or target (information-receiving) hubs. Notable instances of information-receiving hubs include: node 49 of day 16 of culture 1-3, Node 42 of day 22 of culture 2-5 and node 5 of day 15 of culture 2-2 (see Fig. 2b for the node numbers used here). Notable examples of information transmitting hubs include node 28 of day 10 culture 1-3 and nodes 18, 19, 22 and 30 of day 22 of culture 2-5. The specialist computational roles that nodes can take on will be studied in more detail in Sec. II D, with a particular focus on how this relates to the burst propagation.

It is possible to observe some notable instances whereby the information processing properties of a node remain remarkably similar across recording days. For example, nodes 55, 50 and 39 of culture 2-2 are outgoing hubs (with almost no incoming TE) on all 4 recording days. This offers us a tantalising hint that the information processing structure of these networks might be locked in early in development, being reinforced as time progresses. The following subsection (Sec. II C) performs a quantitative analysis of this hypothesis.

### C. Early lock-in of information flows

In the previous subsection, analysis of the functional networks of information flow suggested that the structure of the information processing capacity of the developing networks might be determined early in development and reinforced during the subsequent neuronal maturation.

In order to quantitatively investigate this hypothesis, we examine the relationships in the information flow from a given source to a given target between different recording days. That is, we are probing whether the amount of information flowing between a source and a target on an early day of development will be correlated with the amount flowing on a later day of development. This is equivalent to studying the correlation in the weights of the edges of the functional networks across different recording days. Fig. 3 shows scatter plots between the TE values estimated between each source-target pair on earlier and later days. By observing the pair scatters in Fig. 3a through Fig. 3d we see that, in many pairs of days, there appears to be a substantial correlation between the TE values on the edges across days. This is particularly pronounced for cultures 1-3 and 2-2, though visual assessment of the trend is complicated by the many zero values (where TE is not signficant), gaps in the distribution and outliers. As such, Fig. 3a through Fig. 3d also display the Spearman rank-order correlation (*ρ*) for each early-late pair of days for each culture. This correlation is positive and statistically significant at the *p <* 0.01 level (after Bonferroni correction for multiple comparisons) in 14 out of the 16 early-late pairs of days studied, with the only exceptions being correlations involving the early day 10 for culture 2-5. There are no significant negative correlations. This represents a strong tendency for the relatively strong information flows between a given source and target on later days to be associated with the relatively strong information flow between the same source and target on an earlier day of development. Fig. 3e summarises all Spearman correlations between the early and late TE between source-target pairs. We notice a trend whereby the correlation of the TE values seems to be higher between closer days (sample point being closer to the diagonal) and where those days are later in the development of the cultures (sample points being further to the right).

**FIG. 3:**
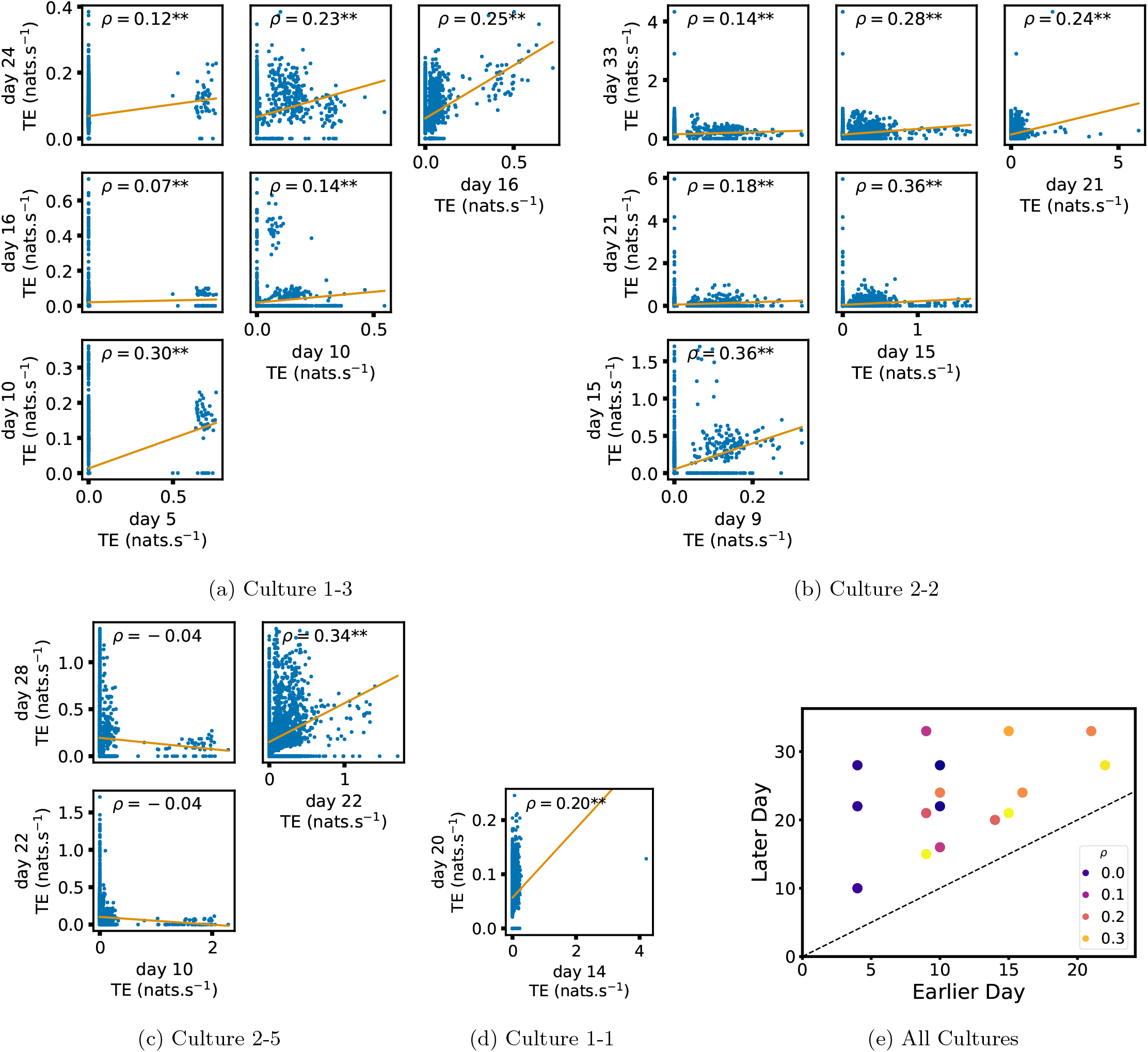
Plots investigating the relationship between the information flow on a given source-target pair over different days of development. (a) through (d) show scatter plots between all pairs of days for each culture (excluding days with zero significant TE values). Specifically, in each scatter plot, the *x* value of a given point is the TE on the associated edge on an earlier day and the *y* value of that same point is the TE on the same edge but on a later day. The days in question are shown on the bottom and sides of the grids of scatter plots. The orange line shows the ordinary least squares regression. The Spearman correlation (*ρ*) between the TE values on the two days is displayed in each plot. Values of *ρ* significant at the 0.05 level are designated with an asterix and those significant at the 0.01 level are designated with a double asterix. A Bonferroni correction for multiple comparisons was used. (e) shows all recording day pairs for all cultures (where the pairs are always from the same culture) and the associated Spearman correlation between the TE on the edges across this pair of recording days.

We also investigated the manner in which a node’s tendency to be an information source hub might be bound early in development. Fig. 4 shows scatter plots between the outgoing TE of each node (averaged across all targets) on different days of development along with the associated Spearman correlations. By observing the scatter plots, it is easy to see that there is a strong positive relationship between the outgoing information flow from a given node on an earlier day of development and the outgoing flow from that same node on a later day. This is not surprising given the correlation we already established for TE on individual pairs, but does not automatically follow from that. More quantitatively, the Spearman correlation between these variables is positive and statistically significant at the *p <* 0.01 level (after Bonferroni correction for multiple comparisons) in 5 out of the 16 early-late pairs of days studied. There is only a single negative correlation and it is not significant. Some of these correlations are particularly strong, and indeed stronger than that observed on the TEs of individual node pairs. For instance, between days 22 and 28 of culture 2-5 we have that *ρ* = 0.69 and between days 9 and 15 of culture 2-2 we have that *ρ* = 0.59. More intriguingly, some of these correlations extend over very large periods of time. Most notably, in culture 2-2, there is a Spearman correlation of *ρ* = 0.52 between the 9th DIV (the first for which there is a recording) and the 33rd DIV. Fig. 4e summarises all Spearman correlations between the early and late total outgoing TE of a given node. As per the TEs for individual node pairs, the correlation is higher between closer days and where those days are later in the development of the cultures.

**FIG. 4:**
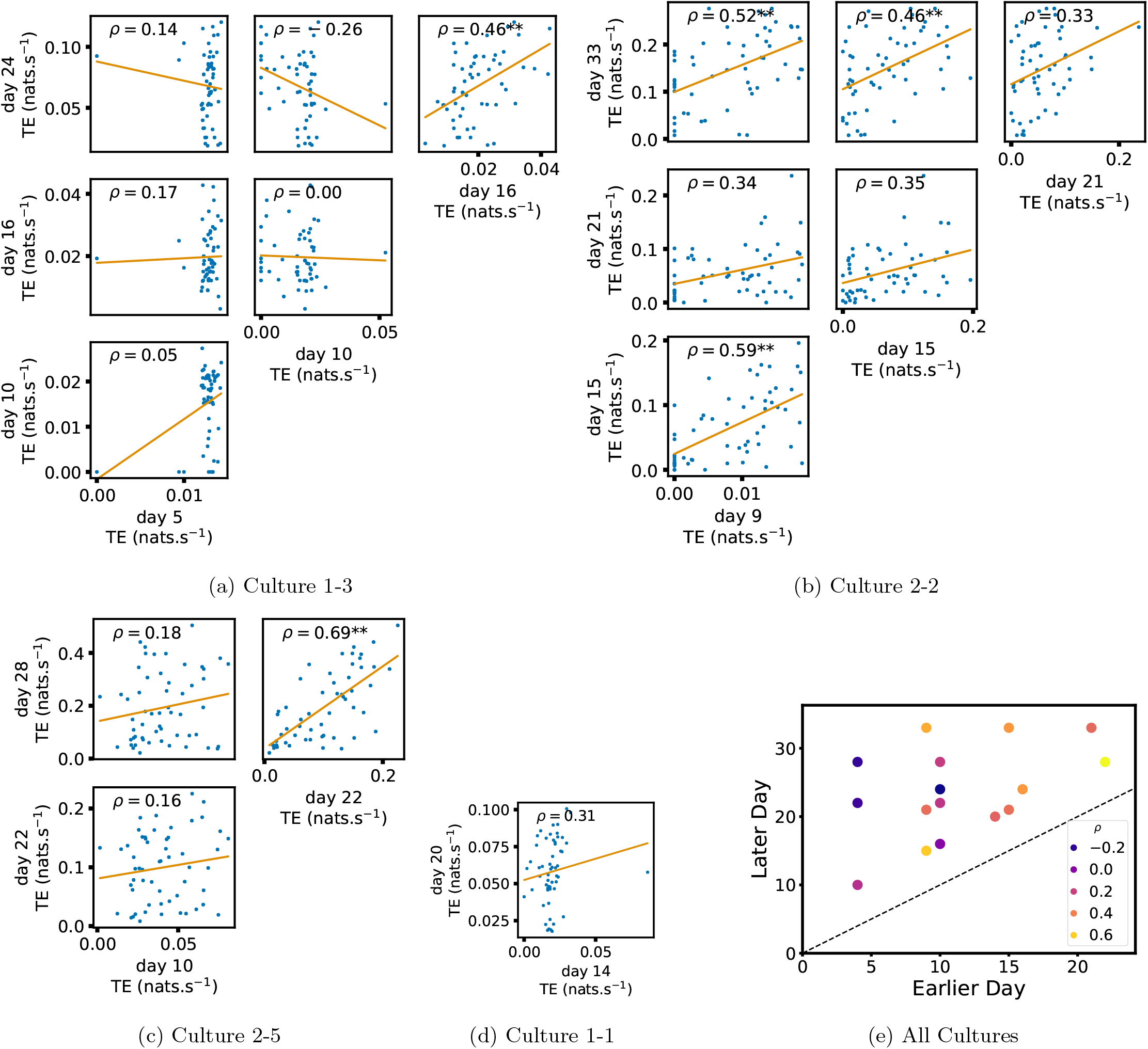
Plots investigating the relationship between the outward information flow from a given node over different days of development. (a) through (d) show scatter plots between all pairs of days for each culture (excluding days with zero significant TE values). Specifically, in each scatter plot, the *x* value of a given point is the average outgoing TE from the associated node on an earlier day and the *y* value of that same point is the total outgoing TE from the same node but on a later day. The days in question are shown on the bottom and sides of the grids of scatter plots. The orange line shows the ordinary least squares regression. The Spearman correlation (*ρ*) between the outgoing TE values on the two days is displayed in each plot. Values of *ρ* significant at the 0.05 level are designated with an asterix and those significant at the 0.01 level are designated with a double asterix. A Bonferroni correction for multiple comparisons was used. (e) shows all recording day pairs for all cultures (where the pairs are always from the same culture) and the associated Spearman correlation between the outward TEs of nodes across this pair of recording days.

Fig. 10 in Appendix B shows similar plots to Fig. 4, but for the average inward TE on each node. As with the average outward TE, in nearly all cases there is a positive correlation between the inward TE on early DIV and the inward TE on later DIV. However, we do observe fewer statistically significant relationships than with the outward TE.

In summary, the data suggests that, in these developing neural cell cultures, the structure of the information flows is to a large degree locked-in early in development. There is a strong tendency for properties of these flows on later days to be correlated with those same properties on earlier days. Specifically, we have looked at the flows between source-target pairs, the average outgoing flow from a source and the average incoming flow to a target. The values of these variables on later DIV were found, in the majority of cases, to be positively correlated with the same values on earlier DIV. Further, there were no cases where a statistically significant negative correlation was found.

### D. Information flows quantify computational role of burst position

Developing cultures of dissociated neurons have a tendency to self-organize so as to produce population bursts or avalanches [21, 25]. Such spike avalanches are not only a feature of cell cultures, being a ubiquitous feature of *in vivo* neural recordings [26–28]. There is a wide body of work discussing the potential computational importance of such periods of neuronal activity [29–35]. It has been observed that cultures often follow an ordered burst propagation [23, 36], whereby some units tend to burst towards the start of the population burst and others tend to burst towards its end. More recent work has proposed that the nodes which burst at different points in this progression play different computational roles [23]. This work has placed special importance on those nodes which burst during the middle of the burst progression, conjecturing that they act as the “brokers of neuronal communication”.

The framework of information dynamics is uniquely poised to illuminate the computational dynamics during population bursting as well as the different roles that might be played by various nodes during these bursts. This is due to its ability to analyse information processing *locally in time* [2, 37–39], as well as directionally between information sources and targets via the asymmetry of transfer entropy. This allows us to isolate the information processing taking place during population bursting activity. We can then determine the information processing roles undertaken by the different nodes and examine how this relates to their position in the burst propagation.

We analyse the information flowing between nodes during population bursts by estimating the *burst-local* TE between nodes in each recording (i.e. averaging the transfer entropy rates only during bursting periods, using probability distribution functions estimated over the whole recordings; see Sec. IV H). We also measure the mean position of each node within bursts (with earlier bursting nodes having a lower numerical position; see Sec. IV G). Fig. 5a and Fig. 5b show plots of the mean burst position plotted against the total inward (Fig. 5a) and outward (Fig. 5b) burst-local TE of each node. Plots are only shown for days where there was a non-zero number of statistically significant burst-local TE values. The Spearman correlation (ρ) between these variables is also displayed on the plots.

**FIG. 5:**
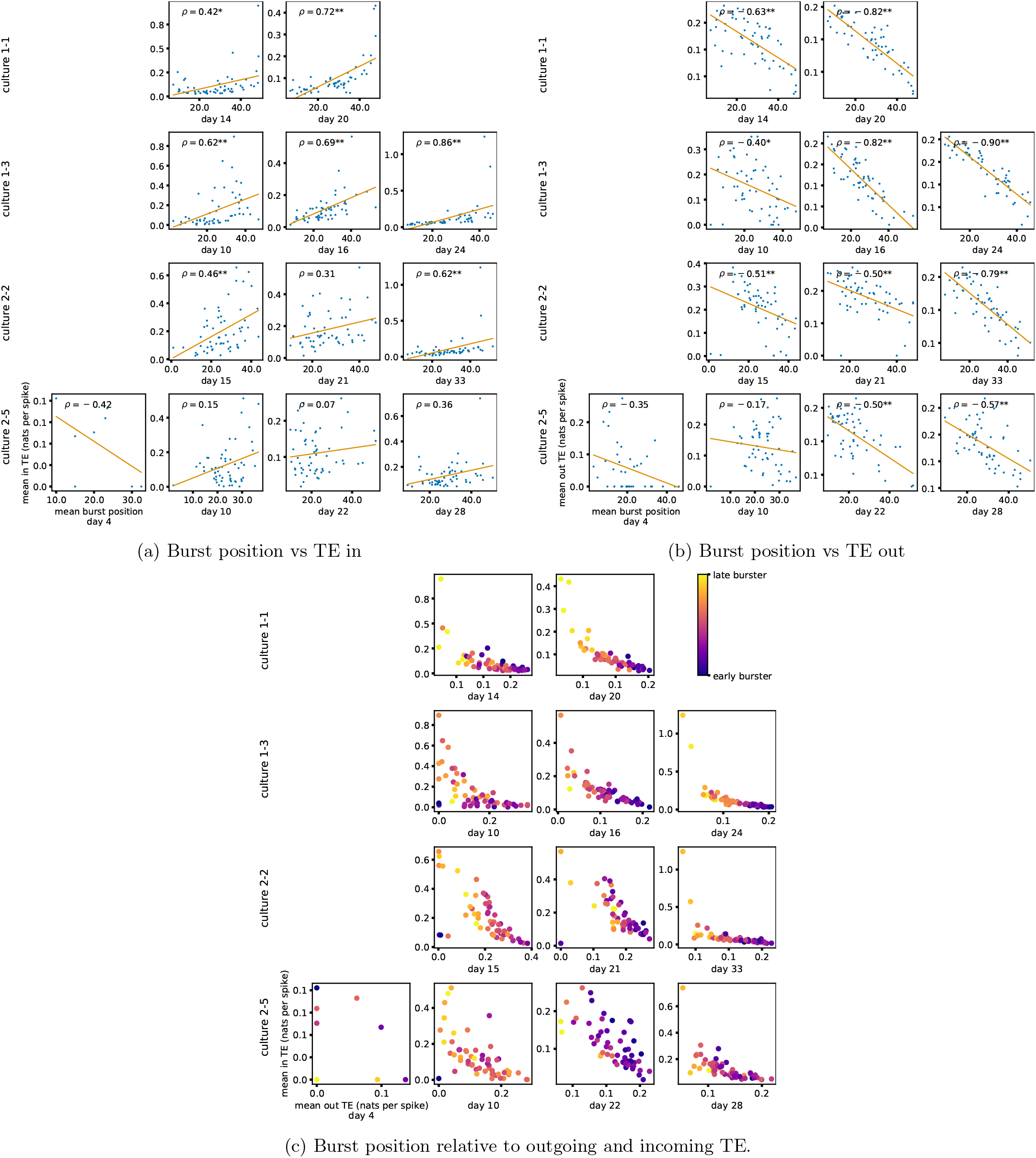
The relationship between the amount of incoming and outgoing local (in burst) TE on a given node and its average burst position. (a) and (b) show the burst position of each node on the *x* axis of each plot, plotted against either the total incoming (a) or outgoing (b) TE on the node. The Spearman correlation (*ρ*) between the mean burst position and the incoming or outgoing TE values is displayed in each plot. Values of *ρ* significant at the 0.05 level are designated with an asterix and those significant at the 0.01 level are designated with a double asterix. A Bonferroni correction for multiple comparisons was used. (c) plots the outgoing TE on the *x* axis and the incoming TE on the *y* axis with the points coloured according to the mean burst position of the node: late bursters are coloured yellow and early bursters are purple.

We see from Fig. 5a that on all days of all cultures (apart from the first recording day of culture 2-5) there is a positive correlation between the mean burst position of the node and the total inward burst-local TE. In other words: later bursting nodes have higher incoming information flows. These correlations are statistically significant at the *p <* 0.01 level (after Bonferroni correction for multiple comparisons) in 6 of the 12 days for which there was a non-zero number of significant burst-local TE values (with 5 of the 12 having *ρ >* 0.5). There are no significant negative correlations. Moreover, the correlations are significant and strong on 3 out of the 4 final days of development, and several are very strong (e.g. the last recording day of culture 1-3 has *ρ* = 0.86). These relationships suggest that there is a tendency for the late bursters to occupy the specialised computational role of information receivers.

Conversely, as shown in Fig. 5b, there is a tendency towards a negative correlation between the mean burst position and the outgoing burst-local TE. On all 12 of the recordings for which there is a non-zero number of significant burst-local TE values we observe a negative Spearman correlation. These correlations are statistically significant at the *p <* 0.01 level (after Bonferroni correction for multiple comparisons) in 9 of the 12 days. More importantly, all values of *ρ* on the final recording day of each culture are significant, with *ρ <* −0.5. These results indicate that the nodes which burst early in the burst propagation tend to occupy the specialised computational role of information transmitters, during the burst period.

Fig. 5c plots the total incoming burst-local TE on each node against the total outgoing burst-local TE, with points coloured according to the node’s mean burst position. We see a very clear pattern in these plots, which is remarkably clear on later recording days: nodes at the beginning of the burst progression have high outgoing information flows with lower incoming flows whereas those at the end of the progression have high incoming flows with lower outgoing flows. By contrast, those nodes at the middle of the burst progression have a balance between outgoing and incoming information transfer. These nodes within the middle of the burst propagation are, therefore, occupying the suggested role of mediators of information flow.

### E. Early lock-in of specialised computational roles

Given that we have seen in Sec. II D that nodes tend to occupy specialised computational roles based on their position in the burst propagation and that we have seen in Sec. II C that information processing properties can lockin early in development, it is worth asking whether the specialised computational roles that nodes occupy during population bursts lock in during the earlier stages of neuronal development.

In order to investigate this question we quantified the computational role occupied by a node by measuring the proportion of its total incoming and outgoing burst-local TE that was made up by its outgoing burst-local TE. These proportions are plotted in Fig. 6 for the different cultures and development days. In order to help us ascertain the relationship over time in these proportions, Fig. 6 shows scatters of these values between earlier and later DIV. It also displays the Spearman rank-order correlations (*ρ*) between the values on different days. Days on which there were no significant burst-local TE values estimated were excluded. On every single pair of days examined, there was a positive Spearman correlation between the proportion of outgoing burst-local TE on the earlier day and this same proportion on the later day. These positive correlations are statistically significant at the *p <* 0.05 level (after Bonferroni correction for multiple comparisons) in 6 out of the 16 early-late pairs of days studied. Fig. 6e summarises all these Spearman correlations between the early and late day pairs.

**FIG. 6:**
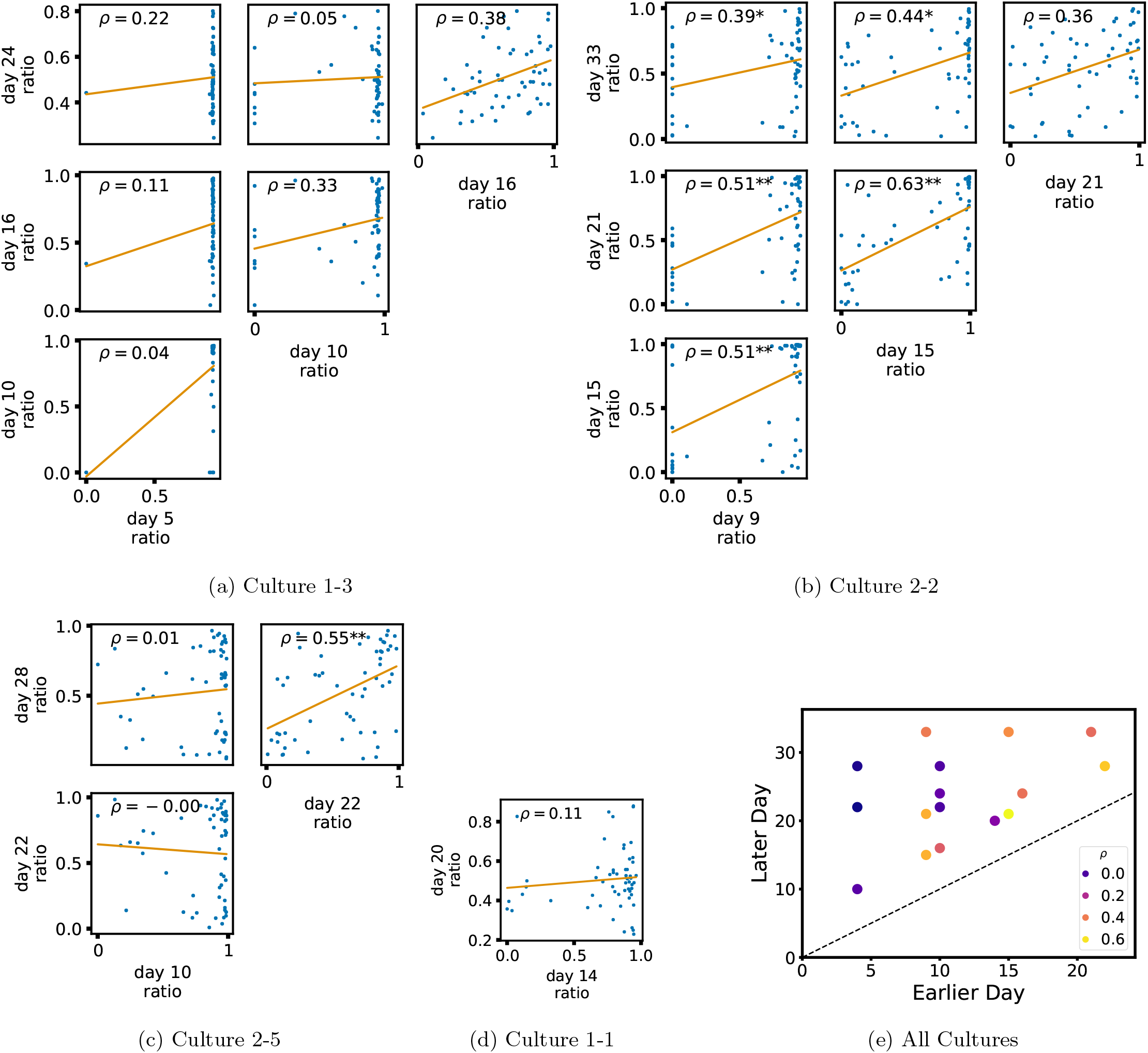
Plots investigating the relationship between the ratio of outward to total burst-local information flow from a given node over different days of development. (a) through (d) show scatter plots between all pairs of days for each culture (excluding days with zero significant burst-local TE values). Specifically, in each scatter plot, the *x* value of a given point is the ratio of total outgoing burst-local TE on the associated node to the total burst-local TE on the same node on one day and the *y* value of that same point is this same ratio on the same node but on a different day. The days in question are shown on the bottom and sides of the grids of scatter plots. The orange line shows the ordinary least squares regression. The Spearman correlation (*ρ*) between the TE values on the two days is displayed in each plot. Values of *ρ* significant at the 0.05 level are designated with an asterix and those significant at the 0.01 level are designated with a double asterix. A Bonferroni correction for multiple comparisons was used. (e) shows all recording day pairs for all cultures (where the pairs are always from the same culture) and the associated Spearman correlation between the outward TE of the nodes across this pair of recording days.

These results suggest that, if a node is an information transmitter during population bursts early in development, it has a tendency to maintain this specialised role later in development. Similarly, being an information receiver early in development increases the probability that the node will occupy this same role later.

### F. Information Flows in an STDP Model of Development

In order to investigate the generality of the phenomena revealed in this paper, we re-implemented a model network [40] of Izhikevich neurons [41] developing according to an STDP [42] update rule as described in Sec. IV B. For the low value of the synaptic time constant which we used (see Sec. IV B), these networks developed from a state where each neuron underwent independent tonic spiking at a regular firing rate, to one in which the dynamics were dominated by periodic population bursts [43]. Small modifications were made to the original model in order that the development occurred over a greater length of time. The greater length of development allowed us to extract time windows which were short relative to the timescale of development (resulting in the dynamics being approximately stationary in these windows) yet still long enough to sample enough spikes for reliable transfer entropy rate estimation. The windows which we used resulted in a median of 5170 spikes per neuron per window, compared with a median of 17 399 spikes per electrode in the biological data. See Sec. IV B for more details on the modifications made. Three windows were extracted, extending between the simulation time-points of 200 and 250 seconds, 400 and 450 seconds and 500 and 550 seconds. These time windows were labelled ‘early’, ‘mid’ and ‘late’, respectively. The early window was chosen such that it had a non-zero number of significant TE values, but such that this number was of the same (order of magnitude in) proportion as observed in the first recording days of the cell cultures (refer to Table II). The mid period was set at the point where population bursting begun to emerge and the late period was set at the point where all neurons were bursting synchronously in a pronounced manner.

TE values between all pairs of model neurons were estimated, as described in Sec. IV D. These estimates were then subjected to the same statistical analysis as the cell culture data, the results of which are presented in the preceding subsections of this Results section. The plots of this analysis are displayed in Fig. 7 and Fig. 8.

**FIG. 7:**
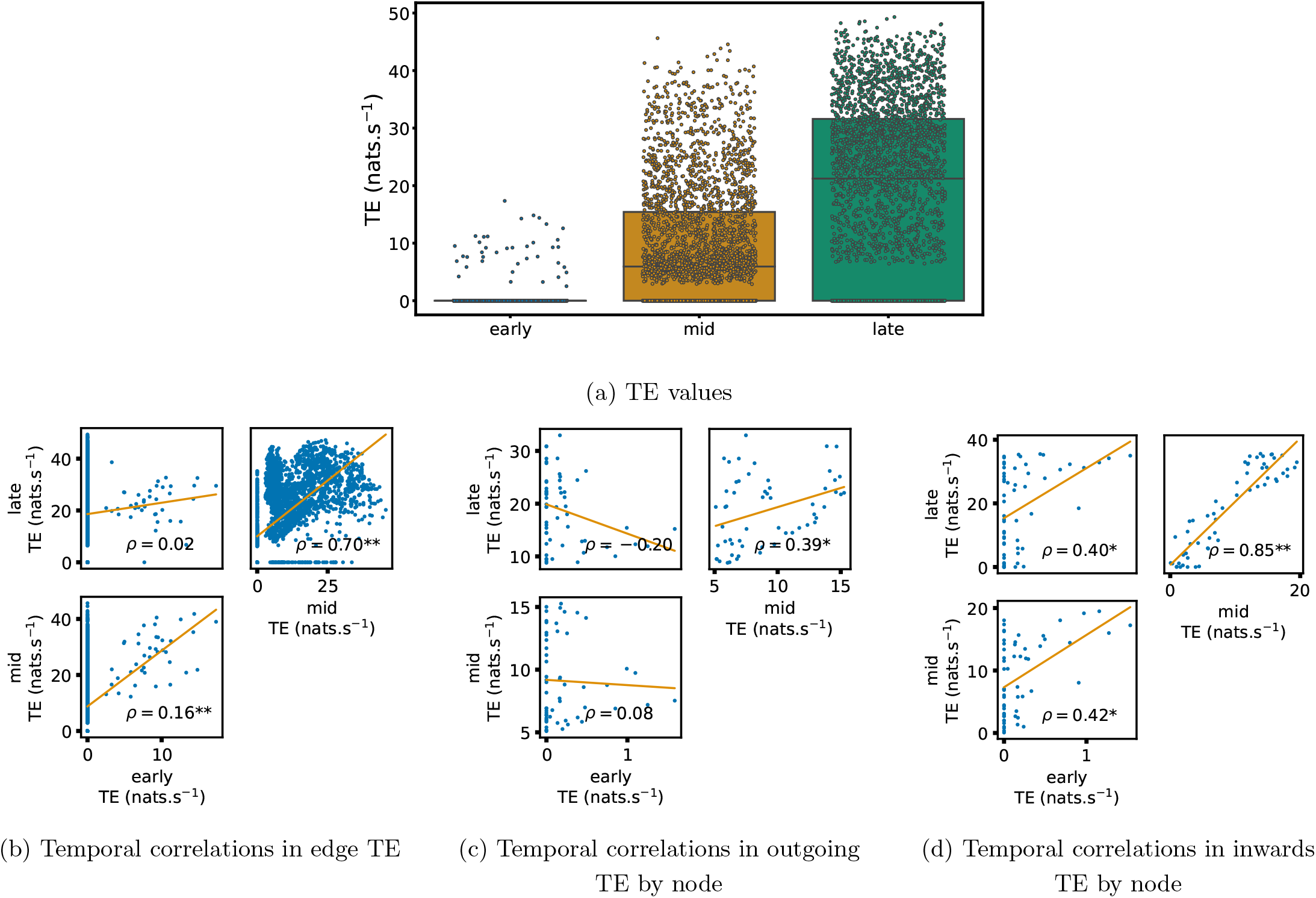
Equivalent plots to those shown in Figs.1, 3, 4 and 10, but for the simulated spiking network developing under STDP. (a) Shows scatters of the TE values overlaid on box plots. The box plots show the quartiles and the median (values greater than 10 standard deviations from the mean have been removed from both the box and scatter plots as outliers). It corresponds to Fig. 1a. (b) through (d) show scatter plots investigating the relationship between TE values (or derived summary statistics) over different stages of development. Specifically, in each scatter plot, the *x* value of a given point is a TE value or derived statistic at an earlier simulation stage and the *y* value of that same point is a TE value (or derived statistic) on the corresponding edge or node, but later in the simulation. The orange line shows the ordinary least squares regression. The Spearman correlation (*ρ*) between the TE values on the two days is displayed in each plot. Values of *ρ* significant at the 0.05 level are designated with an asterix and those significant at the 0.01 level are designated with a double asterix. A Bonferroni correction for multiple comparisons was used. (b) corresponds to the scatter plots in Fig. 3, (c) correponds to the scatter plots in Fig. 4 and (d) correponds to the scatter plots in Fig. 10.

**FIG. 8:**
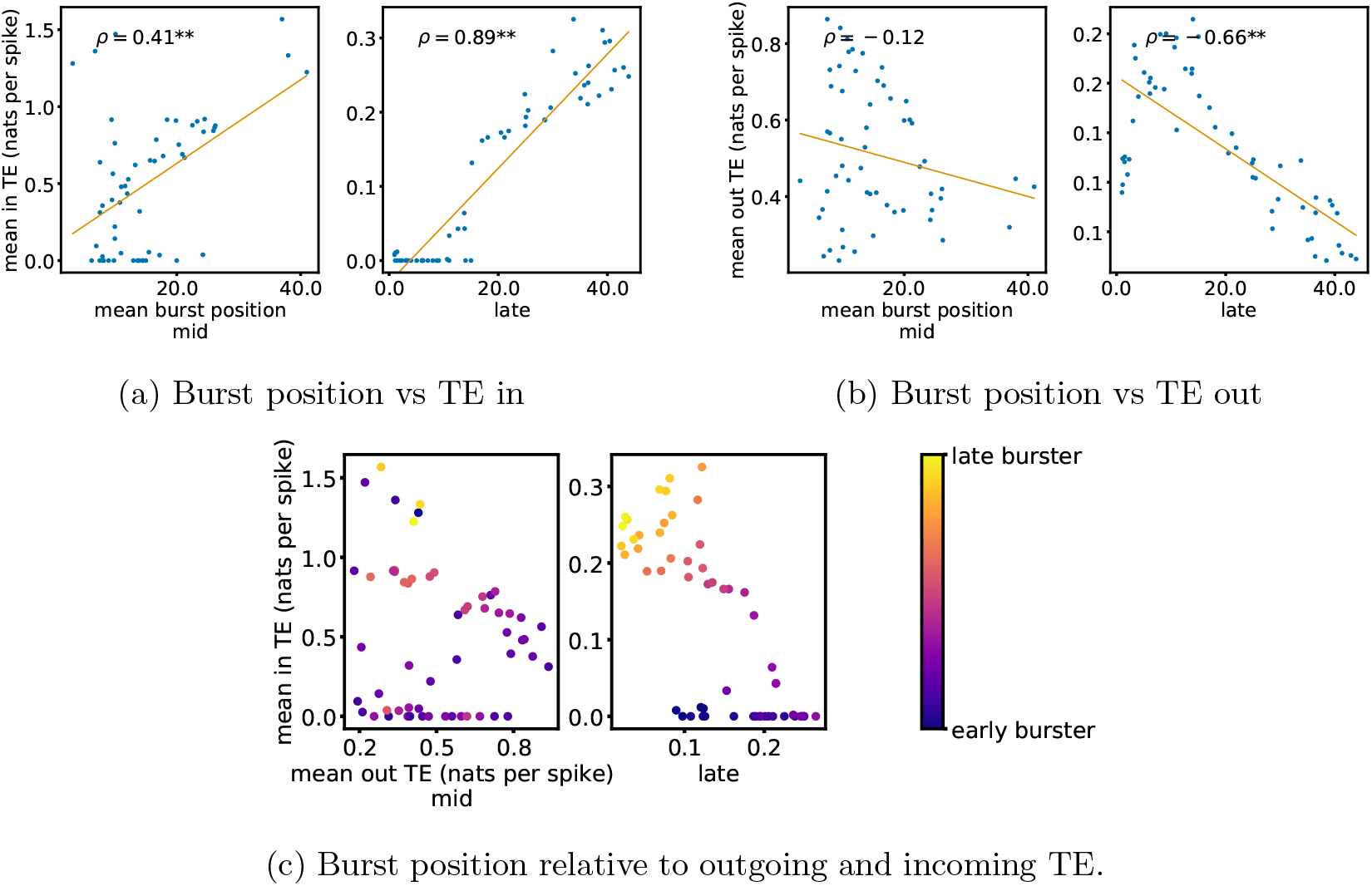
Equivalent plots to those shown in Fig. 5, but for the simulated spiking network developing under STDP. Plots show the relationship between the amount of incoming and outgoing local (in burst) TE on a given node and its average burst position. (a) and (b) show the burst position of each node on the *x* axis of each plot, plotted against either (a) the total incoming or (b) outgoing TE on the node. The Spearman correlation (*ρ*) between the mean burst position and the incoming or outgoing TE values is displayed in each plot. Values of *ρ* significant at the 0.05 level are designated with an asterix and those significant at the 0.01 level are designated with a double asterix. A Bonferroni correction for multiple comparisons was used. (c) Plots the outgoing TE on the *x* axis and the incoming TE on the *y* axis with the points coloured according to the mean burst position of the node: late bursters are coloured yellow and early bursters are purple.

Scatters and box plots of the TE values estimated in each developmental window are shown in Fig. 7a. We observe a large, monotonic, increase in these values over development. This mirrors the finding in cell cultures, as described in Sec. II A.

We also observe the same lock-in phenomenon of information processing as was found in the cell cultures (described in Sec. II C). Fig. 7b through Fig. 7c show the correlation in information flow between different stages of development. Specifically, Fig. 7b shows the correlation in TE values between each ordered pair of neurons between early and later windows. Fig. 7d shows this same correlation, but for the total incoming TE on each neuron and Fig. 7c does this for the total outgoing TE. In six of the nine plotted relationships, we observe a statistically significant positive correlation between values on earlier and later days (significant at the *p <* 0.05 level, with Bonferroni correction). There are no significant negative correlations. As with the cell cultures, some of the observed correlations are particularly strong, such as the Spearman correlation of *ρ* = 0.85 between the total incoming TE on each in the mid window and this same value in the late window. This implies that the spatial structure of the information flow has a tendency to be determined in the earlier stages of development, after which they are locked in – in a similar fashion to what was observed in the biological experiments in earlier sections.

We also performed the same analysis on computational roles as presented in Sec. II D. This analysis, the results of which are presented in Fig. 8, only looked at the mid and late windows. The early window was ignored due to its lack of bursting activity. In the mid recording window, we observe a somewhat weak relationship between the mean burst position of the neuron and its computational role. Fig. 8a shows that there is a weakly significant (at the *p <* 0.01 level) positive correlation between the mean burst position of a neuron and its total incoming burst-local TE (see Sec. IV H for more details on the burst-local TE). There is also a weak negative correlation between the mean burst position and the total outgoing burst-local TE, as shown in Fig. 8a. However, this relationship is not significant. These same figures also display these relationships for the late window. Here, we observe the same directions of relationships, however, they are much stronger and statistically significant in both cases. This implies that we are observing the same specialisation into computational roles based on burst position as was observed in the cell cultures: early bursters display a tendency to be information transmitters, late bursters operate as receivers and middle bursters exhibit a balance of the two.

It is worth noting that the estimated TE values in the model are substantially higher than in the biological dataset. The median estimated TE in the late window of the model was around 20 nats.s^−1^ (Fig. 7a). Conversely, it was less than 0.1 nats.s^−1^ for every last recording day of the cell cultures (Fig. 1a). This is due to the much higher spike rate of the model implying that the dynamics are operating on different time-scales. Indeed, if we compare the magnitude of the burst-local TE — which is measured in nats per spike (see Sec. IV H) — between the model and the biological data (Fig. 8 and Fig. 5, respectively), we find values of similar magnitude.

In summary, in a network model of Izhikevich neurons developing according to STDP towards a state of population bursts, we observe the same developmental information-processing phenomena as in the cell cultures. Namely, the amount of information flowing across the network increases dramatically, the spatial structure of this flow locks in early and the neurons take on specialised computational roles based on their burst position.

## III. DISCUSSION

Biological neural networks are imbued with an incredible capacity for computation, which is deployed in a flexible manner in order to achieve required tasks. Despite the importance of this capacity to the function of organisms, how it emerges during development has remained largely a mystery. Information dynamics [1, 2, 37, 44, 45] provides a framework for studying such computational capacity, by measuring the degree to which the fundamental information processing operations of information storage, transfer and modification occur within an observed system.

Previous work on the information flow component of computational capacity in neural cell cultures [14–20] has focussed on the static structure of information flow networks at single points in time. This has mostly taken the form of elucidating properties of the functional networks implied by the information flows. However, such work leaves open questions concerning how these structures are formed. We address this gap here.

An initial goal in addressing how computational capacity emerges was to determine when the information flow component arrived. It is plausible that this capacity could have been present shortly after plating or that it could have arrived suddenly at a later point in maturation. What we see, however, is that the capacity for information transmission is either not present, or only minimally present, in the early DIV. This can be seen by looking at the very low mean TE values in the first column of Table I. However, over the course of development we see that the TE values increase progressively, reaching values orders of magnitude larger. This implies that information transmission is a capacity which is developed enormously during neuronal development and that its gain is spread consistently throughout the observed period.

The information processing operations of a system tend to be distributed over it in a heterogeneous fashion. For example, it has been found in models of whole-brain networks [46–48], abstract network models [49–51] and even energy networks [52], that nodes with high indegrees tend to also have high outgoing information flows. Sec. II B examined the emergent information flow networks, formed by connecting nodes with a statistically significant TE value between them. In accordance with this previous work – and indeed the large variation in shared, unique and synergistic information flow components observed on the same data set (albeit with the discrete-time estimator) [20] – these networks exhibited a high degree of heterogeneity. Notably, as shown in Fig. 2a, they have prominent hubs of inward flow (sinks) along with less pronounced hubs of outgoing flow (sources). Moreover, along with heterogeneity within individual networks, large structural differences are easily observed between the different networks shown in Fig. 2a.

Keeping with our goal of uncovering how features of mature information flow networks self-organise, we examined how this heterogeneity at both the intra-network and inter-network levels emerged. It was found in Sec. II C that key features of the information flow structure are locked-in early in development. This effect was identified for the outgoing TE from each node for example, where we found strong correlations over the different days of development. It is worth further noting that this lock-in phenomenon occurs remarkably early in development. Specifically, in very many cases, we observe strong correlations between quantities estimated on the first recording days with nonzero TE and the same same quantities estimated on later days. This early lock-in provides us with a mechanism for how the high heterogeneity exhibited in the inflow and outflow hubs emerges. Small differences between networks on early DIV will be magnified on subsequent days. This leads to the high levels of inter-network heterogeneity that we observe. A similar phenomenon has been observed with STDP, which can lead to symmetry breaking in network structure [53, 54], whereby small fluctuations in early development can set the trajectory of the synaptic weights on a specific path with a strong history dependence. In order to confirm a hypothesis that this observed lock-in of information flows could be induced by STDP, in Sec. II F we studied the information dynamics of a model network of Izhikevich neurons developing according to an STDP [42] update rule from a state of independent tonic firing to population bursting. The lock-in of key features of the information flow structure was evident over the period where the network developed from independent firing to synchronous bursting. This indicates a plausible mechanism for our observations, and suggests a broader generality of these phenomena. An interesting difference between the results for the model and the biological data, is that the lock-in was stronger for outward TE in the biological data, whereas it was stronger for inward TE in the model. The reasons for this difference require further investigation, however it might be due to the multi-unit nature of the biological data or the simplicity of the model used.

It has been hypothesised that different neural units take on specialised computational roles [23, 55, 56]. In Sec. II D, we investigated the information flows occurring during the critical bursting periods of the cultures’ dynamics. Specifically, we studied the burst-local TE in order to measure the information being transferred between nodes during these periods. The plots shown in Fig. 5 show a clear tendency for the nodes to take on specialised computational roles, especially later in development. Moreover, these computational roles were tightly coupled to the node’s position in the burst propagation. Nodes initiating the bursts had a tendency to have high outgoing information transfer combined with low incoming information flow, implying their role as information transmitters. The opposite relationship is oberved for late bursters, indicating their role as information receivers. By contrast, nodes bursting during the middle of the progression have a balance between outward and inward flows. This indicates that they are the crucial links between the transmitters and receivers of information. It is worth reflecting on the fact that the observed correlations between burst-local information transfer and burst position will not occur in all bursty neuronal populations. For instance, in populations with periodic bursts, each node’s behaviour will be well explained by its own history, resulting in very low burst-local TE’s, regardless of burst position. Neurons bursting in the middle of the burst progression of dissociated cell cultures have received special attention in past work using undirected measures, where it was conjectured that they act as the “brokers of neuronal communication” [23]. In this work, we have provided novel supporting evidence for this conjecture, by specifically identifying the *directed* information flows into and out of these nodes. Moreover, in Sec. II F, we observed that this same specialisation of neurons into computational roles based on burst position occurred in a model network of Izhikevich neurons which had developed via an STDP learning rule to a state of population bursting. This suggests that this phenomenon might exist more generally than the specific cell cultures studied. It is also worth noting that some of these relationships, notably those shown in Fig. 7b and Fig. 7d are much stronger than what was observed in the cell culture. It is likely that this is due to the fact that in the model we estimated TE between individual model neurons, whereas in the cultures we estimated TE between the multi-unit activity on each electrode.

Returning once more to our focus on investigating the emergence of information flows, we have demonstrated, in Sec. II E, that these specialist computational roles have a tendency to lock in early. There we looked at the ratio of outgoing burst-local TE to the total burst-local TE on each node. It was found that there is a strong tendency for this ratio to be correlated between early and late days of development. This suggests that the computational role that a node performs during population bursts is determined to a large degree early in development.

Insights into development aside, a fundamental technical difference between the work presented here and previous studies of TE in neural cultures is that here we use a recently-developed continuous-time estimator of TE [22]. This estimator was demonstrated to have far higher accuracy in estimating information flows than the traditional discrete-time estimator. The principle challenge which is faced when using the discrete-time estimator is that the curse of dimensionality limits the number of previous time bins that can be used to estimate the history-dependent spike rates. All applications of this estimator to spiking data from cell cultures of which the authors are aware [14–19] made use of only a single previous bin in the estimation of these rates. This makes it impossible to simultaneously achieve high time-precision and capture the dependence of the spike rate on spikes occurring further back in time. Conversely, by operating on the inter-spike intervals, the continuous-time estimator can capture the dependence of the spike rate on events occurring relatively far back in time, whilst maintaining the time precision of the raw data. Looking at a specific representative example, our target history embeddings made use of the previous four inter-spike intervals (Sec. IV E). For the recording on day 24 of culture 1-3, the mean interspike interval was 0.71 seconds. This implies that the target history embeddings on average extended over a period of 2.84 s. The raw data was collected with a sampling rate of 25 kHz [21]. In order to lose no time precision, the discrete-time estimator would thus have to use bins of 40μs, and then in order to extend over 2.84 s, the target history embeddings would therefore need to consist of around 70 000 bins.

It is worth noting that, as we were performing a longitudinal analysis where each studied recording was separated by days or weeks, we did not perform spike sorting as we would have been unable to match the different units on an electrode across different recordings. We would then not have been able to compare the TE values on a given unit over the course of development. Instead, we analyzed the spikes on each electrode without sorting. As such, this work studies multi-unit activity [23]. Spike sorting applied to data collected from a near-identical recording setup found an average of four neurons per electrode [57]. This situates this work at a spatial scale slightly larger than spike-sorted neural data, but still orders of magnitude finer than fMRI, EEG or MEG [58].

An exciting direction for future work will be to examine the information flow provided by higher-order multivariate TEs [59, 60]. The networks inferred by such higher-order TEs are able to better reflect the networks’ underlying structural features [59]. As was the case with bivariate TEs prior to this work, there is an absence of work investigating how the networks of multivariate information flow emerge during neural development. Moreover, moving to higher-order measures will allow us to more fully characterise the multifaceted specialised computational roles undertaken by neurons.

## IV. METHODS

### A. Cell culture data

The spike train recordings used in this study were collected by Wagenaar et. al. [21] and are freely available online [24]. The details of the methodology used in these recordings can be found in the original publication [21]. A short summary of their methodology follows:

Dissociated cultures of rat cortical neurons had their activity recorded. This was achieved by plating 8×8 Multi-Electrode Arrays (MEAs), operating at a sampling frequency of 25 kHz with neurons obtained from the cortices of rat embryos. The spacing between the electrodes was 200μm center-to-center. The MEAs did not have electrodes on their corners and one electrode was used as ground, resulting in recordings from 59 electrodes. In all recordings, electrodes with less than 100 spikes were removed from the analysis. This resulted in electrodes 37 and 43 (see Fig. 2b for the position of these electrodes) being removed from every recording as no spikes were recorded on them. The spatial layout of the electrodes is available from the website associated with the dataset [24], allowing us to overlay the functional networks onto this spatial layout as is done in figure Fig. 2a.

30 minute recordings were conducted on most days, starting from 3-4 Days *In Vitro* (DIV). The end point of recording varied between 25 and 39 DIV. Longer overnight recordings were also conducted on some cultures at sparser intervals. As the accurate estimation of information-theoretic quantities requires substantial amounts of data [22, 61], in this work we make use of these longer overnight recordings. These recordings were split into multiple files. The specific files used, along with the names of the cultures and days of the recordings are listed in Table III.

**TABLE III:** File numbers used for each culture on each day. These correspond to the file numbering used in the freely available dataset used in this study, provided by Wagenaar et. al.[21, 24]

|  |  |  |  |  |
| --- | --- | --- | --- | --- |
| **Culture 1-1** | day 4 | day 14 | day 20 |  |
|  | 2 | 2 | 2 |  |
| **Culture 1-3** | day 5 | day 10 | day 16 | day 24 |
|  | 2 | 2 | 2 | 2 |
| **Culture 2-2** | day 9 | day 15 | day 21 | day 33 |
|  | 2 | 2 | 2 | 2 |
| **Culture 2-2** | day 4 | day 10 | day 22 | day 28 |
|  | 1 | 1 | 2 | 1 |

The original study plated the electrodes with varying densities of cortical cells. However, overnight recordings were only performed on the ‘dense’ cultures, plated with a density of 2500 cells*/*μL.

The original study performed threshold-based spike detection by determining that a spike was present in the case of an upward or downward excursion beyond 4.5 times the estimated RMS noise of the recorded potential on a given electrode. The analysis presented in this paper makes use of these detected spike times. No spike sorting was performed and, as such, we are studying multi-unit activity (MUA) [23].

### B. Network of Izhikevich Neurons

The model spiking network used to generate the data analysed in Sec. II F is identical to that presented in [40], with a few minor alterations. This model consists of Izhikevich neurons [41] developing according to an STDP [42] update rule. At the beginning of the simulation, each neuron performs independent tonic spiking, however, the network develops towards population bursts.

The specific model settings used were based on those used to produce Fig. 5A in [40]. That is, the proportion of inhibitory neurons (*α*) and the synapse time delay (*τ*_*ij*_) were both set to 0. The first change made was to use 59 neurons, as opposed to the 500 used in [40], in order to correspond to the number of electrodes used in the cell culture recordings. The maximum connection strength (*g*_*max*_) was also increased from 0.6 to 10 in order to compensate for this reduction in the network size.

The only remaining change was made in order to slow the rate of development of the population. The reasoning behind this was to allow for the extraction of windows which were much shorter than the time scale of development, resulting in the dynamics within these windows being approximately stationary (and including enough samples for estimation of the transfer entropy rates). Specifically, this change was to greatly reduce the values of the maximum synaptic potentiation and depression (*A*_+_ and *A*_−_). These values were reduced from 5*×*10^−2^ to 4*×*10^−4^.

### C. Data pre-processing

As the data was sampled at 25 kHz, uniform noise distributed between −20μs and 20μs was added to each spike time. This is to prevent the TE estimator from exploiting the fact that, in the raw data, inter-spike intervals are always an integer multiple of 40μs.

### D. Transfer entropy estimation

The (bivariate) Transfer entropy (TE) [3, 4] was estimated between each pair of electrodes in each of the recordings listed in Table III. TE is the mutual information between the past state of a source process and the present state of a target process, conditioned on the past state of the target. More specifically (in discrete time), the TE rate is:

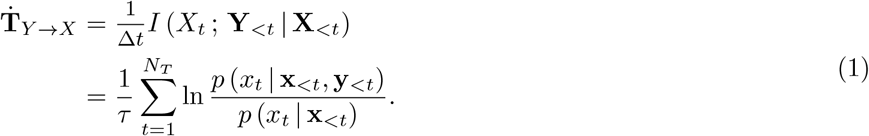

The TE above is being measured from a source *Y* to a target *X, I*(*·* ; *·* | *·*) is the conditional mutual information [62], *x*_*t*_ is the current state of the target, **x**_*<t*_ is the history of the target, **y**_*<t*_ is the history of the source, Δ*t* is the bin width (in time units), *τ* is the length of the processes and *N*_*T*_ = *τ/*Δ*t* is the number of time samples (bins). The histories **x**_*<t*_ and **y**_*<t*_ are usually captured via embedding vectors, e.g. **x**_*<t*_ = **x**_*t*−*m*:*t*−1_ = {*x*_*t*−*m*_, *x*_*t*−*m*+1_, …, *x*_*t*−1_}.

#### 1. Previous application of the discrete-time estimator

Previous applications of TE to spiking data from neural cell cultures [14–20] made use of this discrete-time formulation of TE. This work was primarily focussed on the directed functional networks implied by the estimated TE values between pairs of nodes which has revealed interesting features of the information flow structure. Shimono and Beggs [15] found that these networks exhibited a highly non-random structure and contained a long-tailed degree distribution. This work was expanded by Nigam et. al.[14], where it was found that the functional networks contained a rich-club topology. Conversely, Timme et. al. [17] found that the hubs of these networks were localised to certain time scales. Other work [19, 20] has instead focussed on how the components of information flows in cell cultures can be decomposed into unique, redundant and synergystic components.

#### 2. Continuous-time estimation

It has, relatively recently, been shown that, for event-based data such as spike-trains, in the limit of small bin size, that the TE is given by the following expression [63]:

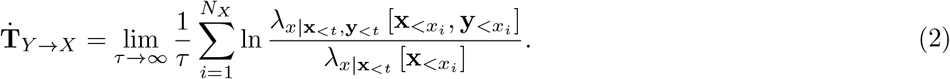

Here, 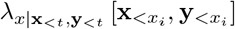 is the instantaneous firing rate of the target conditioned on the histories of the target 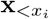 and source 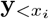 at the time points *x*_*i*_ of the spike events in the target process. 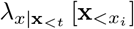 is the instantaneous firing rate of the target conditioned on its history alone, ignoring the history of the source. It is important to note that the sum is being taken over the *N*_*X*_ spikes of the target: thereby evaluating log ratios of the expected spike rates of the target given source and target histories versus target histories alone, *when* the target does spike. As this expression allows us to ignore the “empty space” between events, it presented clear potential for allowing for more efficient estimation of TE on spike trains.

This potential was recently realised in a new continuous-time estimator of TE presented in [22] (and utilised in [64]), and all TE estimates in this paper were performed using this new estimator. In [22] it is demonstrated that this continuous-time estimator is far superior to the traditional discrete-time approach to TE estimation on spike trains. For a start, unlike the discrete-time estimator, it is consistent. That is, in the limit of infinite data, it will converge to the true value of the TE. It was also shown to have much preferable bias and convergence properties. Most significantly, perhaps, this new estimator utilises the inter-spike intervals to efficiently represent the history embeddings 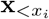 and 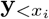 in estimating the relevant conditional spike rates in (2). This then allows for the application of the highly effective nearest-neighbour family of information-theoretic estimators [61, 65], which bring estimation efficiency, bias correction, and together with their application to inter-spike intervals enable capture of long time-scale dependencies. This is in contrast with the traditional discrete-time estimator which uses the presence or absence of spikes in time bins as its history embeddings (it sometimes also uses the number of spikes occurring in a bin). In order to avoid the dimensionality of the estimation problem becoming sufficiently large so as to render estimation infeasible, only a small number of bins can be used in these embeddings. Indeed, to the best of the authors’ knowledge, all previous applications of the discrete-time TE estimator to spiking data from cell cultures used only a single bin in their history embeddings. The bin widths used in those studies were 40μs [14], 0.3 ms [66], and 1 ms [15, 67]. Some studies chose to examine the TE values produced by multiple different bin widths, specifically: 0.6 ms and 100 ms [16], 1.6 ms and 3.5 ms [19] and 10 different widths ranging from 1 ms to 750 ms [17]. And specifically, those studies demonstrated the unfortunate high sensitivity of the discrete-time TE estimator to the bin width parameter. In the instances where narrow (*<* 5 ms) bins were used, only a very narrow slice of history is being considered in the estimation of the history-conditional spike rate. This is problematic, as it is known that correlations in spike trains exhist over distances of (at least) hundreds of milliseconds [68, 69]. Conversely, in the instances where broad (*>* 5 ms) bins were used, relationships occurring on fine time scales will be completely missed. This is significant given that it is established that correlations at the millisecond and sub-millisecond scale play a role in neural function [70–73]. In other words, previous applications of transfer entropy to electrophysiological data from cell cultures either captured some correlations occurring with fine temporal precision or they captured relationships occurring over larger intervals, but never both simultaneously. This can be contrasted with the inter-spike interval history representation used in this study. To take a concrete example, for the recording on day 24 of culture 1-3, the average interspike interval was 0.71 seconds. This implies that the target history embeddings (composed of 4 inter-spike intervals) on average extended over a period of 2.84 s and the source history embeddings (composed of 2 inter-spike intervals) on average extended of a period of 1.42 s. This is despite the fact that our history representations retain the precision of the raw data (40μs) and the ability to measure relationships on this scale where they are relevant (via the underlying nearest-neighbour estimators).

The parameters used with this estimator are shown in Table IV. The values of *k*_global_ and *k*_perm_ were chosen because, in previous work [22], similar values were found to facilitate stable performance of the estimator. The high values of *N*_*U*_ and *N*_*U*,surrogates_ were chosen so that histories during bursting periods could be adequately sampled. These two parameters refer to sample points placed randomly in the spike train, at which history embeddings are sampled. As the periods of bursting comprise a relatively small fraction of the total recording time, many samples need to be placed in order to achieve a good sample of histories potentially observed during these periods. The choice of embedding lengths is discussed in the next subsection (Sec. IV E) and the choice of *N*_surrogates_ is discussed in Sec. IV F.

**TABLE IV:** The parameter values used in the continuous-time TE estimator. A complete description of these parameters, along with analysis and discussion of their effects can be found in [22].

| Parameter | Description | Value |
| --- | --- | --- |
| $N_X$ | Number of spikes in the target spike train | varied (see text) |
| $l_X$ | Number of inter-spike intervals in target history embeddings | 4 |
| $l_Y$ | Number of inter-spike intervals in source history embeddings | 2 |
| $k_{\text{global}}$ | Number of nearest neighbours to find in the initial search | 10 |
| $k_{\text{perm}}$ | Number of nearest neighbours to consider during surrogate generation | 4 |
| $N_U$ | Number of random samples of histories at non-spiking points in time | $10N_X$ |
| $N_{U,\text{surrogates}}$ | Number of random samples of histories at non-spiking points in time used for surrogate generation | $10N_X$ |
| $N_{\text{surrogates}}$ | Number of surrogates to generate for each node pair | 100 |

Instead of selecting a single number of target spikes *N*_*X*_ to include in the analysis, we chose to include all the spikes that occurred within the first hour of recording time. The reason for doing this was that the spike rates varied by orders of magnitude between the electrodes. This meant that fixing the number of target spikes would result in the source spikes being severely undersampled in cases where the target spike rate was much higher than the source spike rate. When using one hour of recording time, the smallest number of spikes per electrode was 481, the maximum was 69627 and the median was 17 399.

### E. Selection of embedding lengths

The target embedding lengths were determined by adapting the technique ([60, 74] extending [75]) of maximising the bias-corrected Active Information Storage (AIS) [45] over different target embedding lengths for a given target. Our adaptations sought to select a consensus embedding parameter for all targets on all trials, to avoid different bias properties due to different parameters across targets and trials, in a similar fashion to [76]. As such, our approach determines a target embedding length *l*_*X*_ which maximises the *average* bias-corrected AIS across all electrodes, using one representative recording (selected as day 23 of culture 1-3). To estimate AIS within the continuous-time framework [63] for this purpose, we estimated the difference between the second KL divergence of eq. (10) of [22] and the mean firing rate of the target. These estimates contain inherent bias-correction, as per the TE estimator itself. Moreover, the mean of surrogate values was subtracted to further reduce the bias. The embedding length *l*_*X*_ was continuously increased so long as each subsequent embedding produced a statistically significant (at the *p <* 0.05 level) increase in the average AIS across the electrodes. The resulting mean AIS values (along with standard deviations) and *p*-values are shown in Table V. We found that every increase in *l*_*X*_ up to 4 produced a statistically significant increase in the mean AIS. The increase from 4 to 5 produced a non-significant decrease in the mean AIS and so *l*_*X*_ was set to 4.

**TABLE V:**
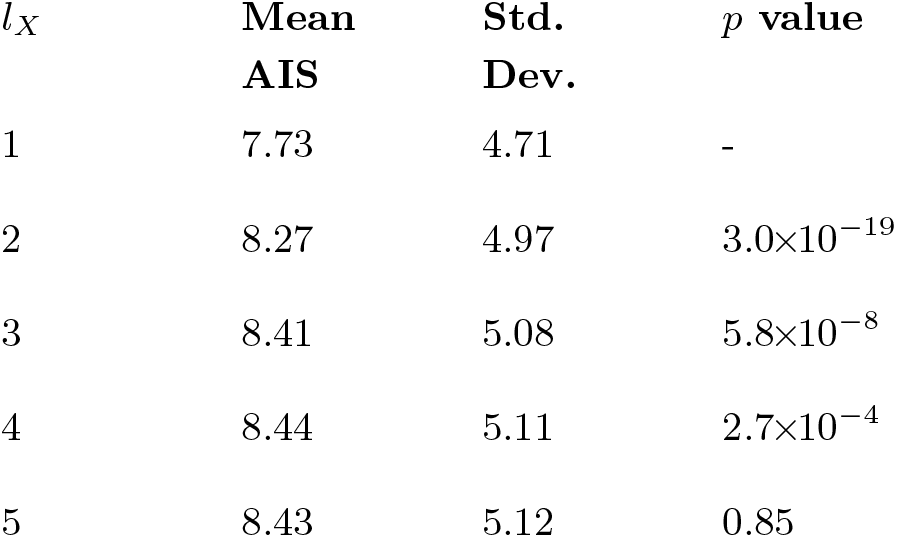
Summary statistics for the AIS values estimated at different target embedding lengths *l*_*X*_ across all electrodes of a representative recording (day 23 of culture 1-3). The *p* values shown in the fourth column are associated with the null hypothesis that the mean AIS at the given *l*_*X*_ is equal to the mean AIS at *l*_*X*_ − 1.

| $l_X$ | Mean<br>AIS | Std.<br>Dev. | $p$ value |
| --- | --- | --- | --- |
| 1 | 7.73 | 4.71 | - |
| 2 | 8.27 | 4.97 | $3.0 \times 10^{-19}$ |
| 3 | 8.41 | 5.08 | $5.8 \times 10^{-8}$ |
| 4 | 8.44 | 5.11 | $2.7 \times 10^{-4}$ |
| 5 | 8.43 | 5.12 | 0.85 |

With the target embedding length determined, we set about similarly determining a consensus source embedding length *l*_*Y*_ by estimating the TE between all directed electrode pairs on the same representative recording for different values of *l*_*Y*_. Each estimate also had the mean of the surrogate population subtracted to reduce its bias (see Sec. IV F).

The embedding length was continuously increased so long as each subsequent embedding produced a statistically significant (at the *p <* 0.05 level) increase in the average TE across all electrode pairs. The resulting mean TE values (along with standard deviations) and *p*-values are shown in Table VI. We found that increasing *l*_*Y*_ from 1 to 2 produced a statistically significant increase in the mean TE. However, increasing *l*_*Y*_ from 2 to 3 produced a non-significant decrease in the mean TE. As such, we set *l*_*Y*_ to 2

**TABLE VI:** Summary statistics for the TE values estimated at different source embedding lengths *l*_*Y*_ between all electrodes of a representative recording (day 23 of culture 1-3). The *p* values shown in the fourth column are associated with the null hypothesis that the mean TE at the given *l*_*Y*_ is equal to the mean TE at *l*_*Y*_ − 1.

| $l_Y$ | Mean<br>TE | Std.<br>Dev. | $p$ value |
| --- | --- | --- | --- |
| 1 | 0.031 | 0.043 | - |
| 2 | 0.058 | 0.056 | 0.0 |
| 3 | 0.057 | 0.069 | 0.84 |

### F. Significance testing of TE values

In constructing the directed functional networks displayed in Fig. 2a, we tested whether the estimated TE between each source-target pair was statistically different from the distribution of TEs under the null hypothesis of conditional independence of the target from the source (i.e. TE consistent with zero). Significance testing for TE in this way is performed by constructing a population of surrogate time-series or history embeddings that conform to the null hypothesis of zero TE [59, 60, 77]. We then estimate the TE on each of these surrogates to generate a null distribution of TE. Specifically, we generate the surrogates and compute their TEs according the method associated with the continuous-time spiking TE estimator [22] and using the parameters shown in Table IV. One small change was made to that surrogate generation method: instead of laying out the *N*_*U*,surrogates_ sample points randomly uniformly, we placed each one at an existing target spike, with the addition of uniform noise on the interval [−80 ms, 80 ms]. This was to ensure that these points adequately sampled the incredibly dense burst regions.

With the surrogate TE distribution constructed, the resulting *p* value for our TE estimate can be computed by counting the proportion of these surrogate TEs that are greater than or equal to the original estimate. Here, we seek to compare significance against a threshold of *α <* 0.01. We chose this lower threshold as false positives are generally considered more damaging than false negatives when applying network inference to neuroscientific data [78]. We also applied a Bonferroni correction [79] to all the significance tests done on a given recording. Given that there are 59 electrodes in the recordings, 3422 tests were performed in each recording. This meant that, once the Bonferroni correction was included, the significance threshold dropped to *p <* 2.9*×*10^−6^. Comparing against such a low significance threshold would require an infeasible number of surrogates for the many pairs within each recording, if computing the *p* value by counting as above. Instead, we assume that the null TE distribution is Gaussian, and compute the *p* value for our TE estimate using the CDF of the Gaussian distribution fitted from 100 surrogates (as per e.g. [7]). Specifically, the *p* value reports the probability that a TE estimate on history embeddings conforming to the null hypothesis of zero TE being greater than or equal to our original estimated TE value. If this *p* value is below the threshold then the null hypothesis is rejected and we conclude that there is a statistically significant information flow between the electrodes.

### G. Analysis of population bursts

A common family of methods for extracting periods of bursting activity from spike-train recordings examines the length of adjacent inter-spike intervals. The period spanned by these intervals is designated a burst if some summary statistic of the intervals (e.g.: their sum or maximum) is below a certain threshold [21, 80–83]. In order to detect single-electrode as well as population-wide bursts, we implement such an approach here.

We first determine the start and end points of the bursts of each individual electrode. The locations of the population bursts were subsequently determined using the results of this per-electrode analysis.

The method for determining the times during which an individual electrode was bursting proceded as follows: The spikes were moved through sequentially. If the interval between a given spike and the second most recent historic spike for that electrode was less than *α*, then, if the electrode was not already in a burst, it was deemed to have a burst starting at the second most recent historic spike. A burst was taken to continue until an inter-spike interval greater than *a* ∗ *α* was encountered. If such an interval was encountered, then the end of the burst was designated as the timestamp of the earlier of the two spikes forming the interval.

The starts and ends of population bursts were similarly determined by moving through the timeseries in a sequential fashion. If the population was not already designated to be in a burst, but the number of electrodes currently bursting was greater than the threshold *β*, then a burst start position was set at the point this threshold was crossed. Conversely, if the electrode was already designated to be in a burst and the number of individual electrodes currently bursting dropped below the threshold *γ* (*γ < β*), then a burst stop position was set at the point this threshold was crossed.

In this paper, we always made use of the parameters *α* = 16 ms, *a* = 3, *β* = 15 and *γ* = 10. These parameters were chosen by trial-and-error combined with visual inspection of the resulting inferred burst positions. The results of this scheme showed low sensitivity to the choice of these parameters.

### H. Estimation of burst-local TE

The information dynamics framework provides us with the unique ability to analyse information processing locally in time [2, 37, 38]. We make use of that ability here to allow us to specifically examine the information flows during the important period of population bursts. The TE estimator which we are employing here [22] sums contributions from each spike in the target spike train. It then divides this total by the time length of the target spike train that is being examined. In order to estimate the burst-local TE, we simply sum the contributions from the target spikes where those spikes occurred during a population burst. We then normalise by the number of such spikes, providing us with a burst-local TE estimate in units of nats per spike, instead of nats per second.

## CONTRIBUTIONS

**David P. Shorten** Designed research, Analyzed data, Performed research, Wrote the paper, Edited the paper

**Michael Wibral** Designed research, Edited the paper

**Viola Priesemann** Designed research, Edited the paper

**Joseph T. Lizier** Designed research, Wrote the paper, Edited the paper, Supervision, Funding Acquisition

## ACKNOWLEDGEMENTS

Joseph Lizier was supported through the Australian Research Council DECRA grant DE160100630 - www.arc.gov.au/grants/discovery-program/discovery-early-career-researcher-award-decra and The University of Sydney Research Accelerator (SOAR) prize program - sydney.edu.au/research/our-researchers/sydney-researchaccelerator html. The authors acknowledge the technical assistance provided by the Sydney Informatics Hub, a Core Research Facility of the University of Sydney. In particular, the analysis presented in this work made use of the Artemis HPC cluster.

## Appendix A: Distribution of information flow values

Previous studies have placed an emphasis on the observation of log-normal distributions of TE values in *in vitro* cultures of neurons [14, 15]. As such, we analysed the distribution of the nonzero (statistically significant) estimated TE values in each individual recording.

Fig. 1b shows histograms as well as probability density functions estimated by a kernel density estimator (KDE) of the nonzero TE values for each recording. From these plots we can see that the distributions of TE values exhibits a clear right (positive) skew.

In order to ascertain how well the estimated TE values were described by a log-normal distribution, we constructed Quantile-Quantile (QQ) plots [84] for the TE values against the log-normal distribution in figure Fig. 9b. In all recordings, the plotted points deviate from the line *y* = *x*, indicating that the data is not well described by a log-normal distribution. However, this deviation appears only slight for some recordings, most notably days 22 and 28 of culture 2-5. We also perform Shapiro-Wilke tests [85] for log-normality, the resulting *p* values are displayed in Table VIII. The *p* values for every recording are incredibly low, meaning that we reject the null hypothesis of a log-normal distribution in every case.

**FIG. 9:**
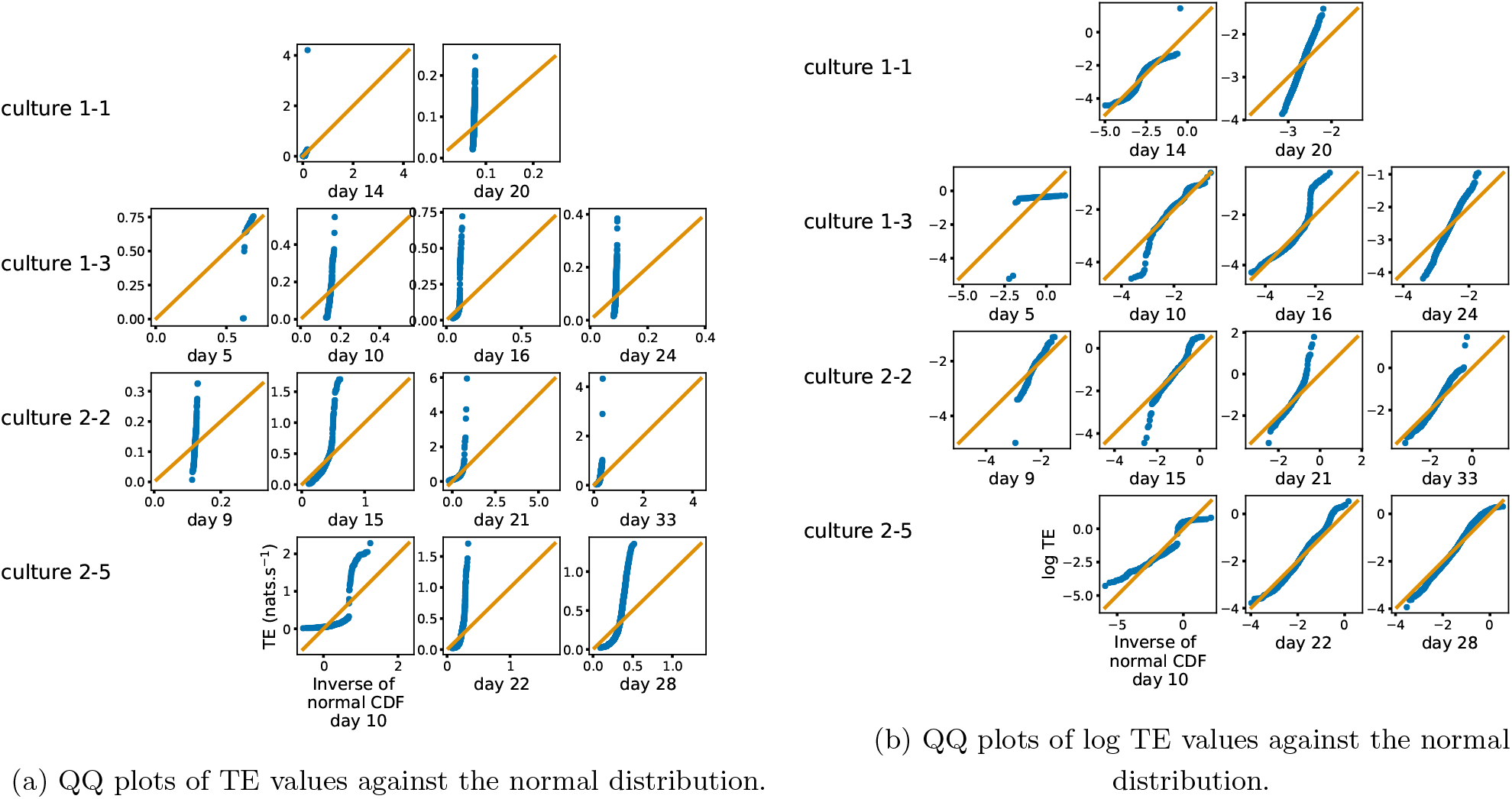
Quantile-Quantile (QQ) plots [84] of the nonzero estimated TE values against normal and log-normal distributions, respectively. The *y* axis shows estimated TE values (or their logarithm) whereas the *x* axis shows the value of the normal distribution at the same quantile. The solid orange line shows the line *y* = *x*. If the data is drawn from the distribution against which it is being plotted then the blue marks will sit along this line. We observe that the distributions of TE values deviate substantially from both normal and log-normal distributions in all recordings analysed.

**FIG. 10:**
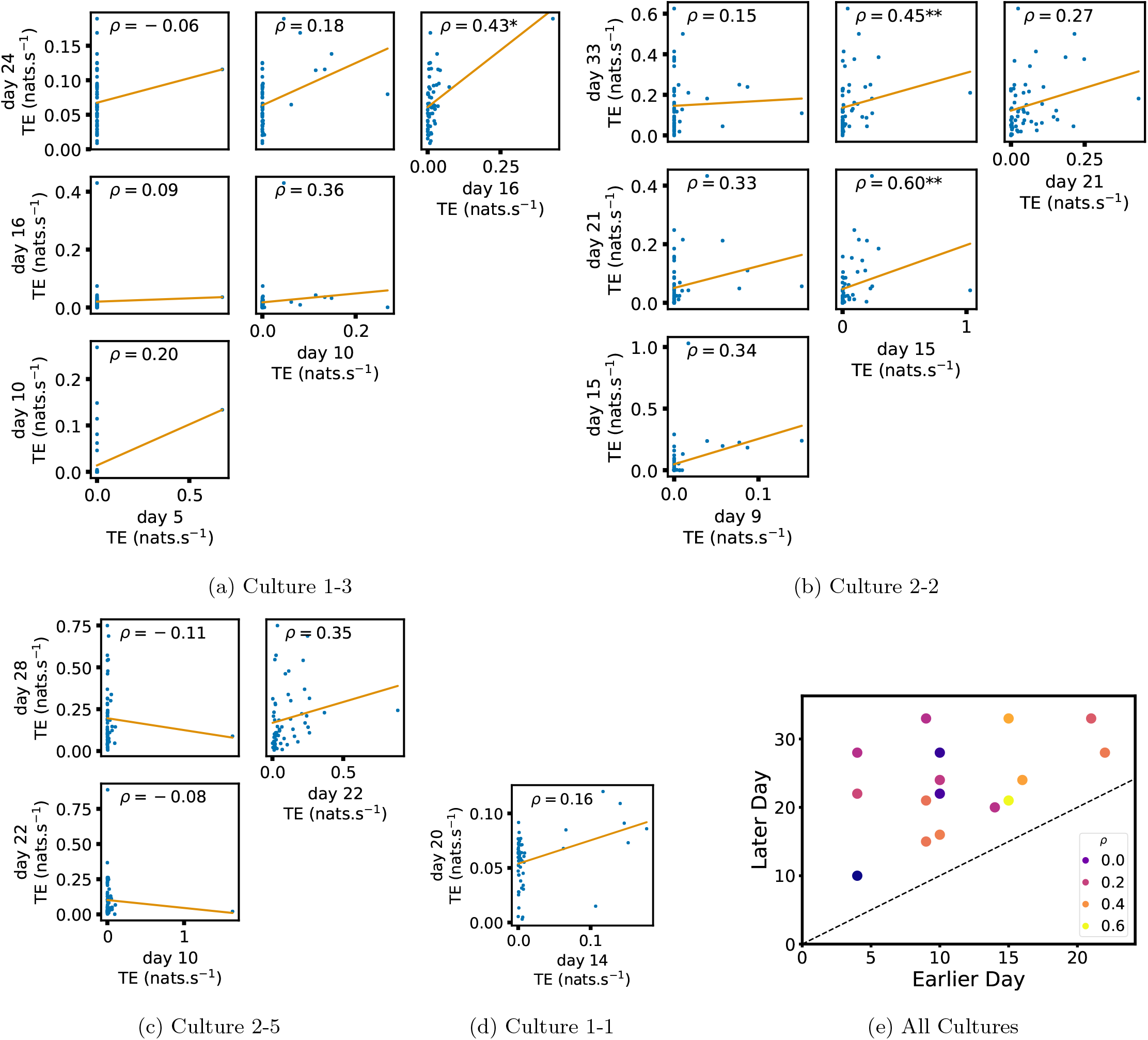
Plots investigating the relationship between the inward information flow from a given node over different days of development. (a) through (d) show scatter plots between all pairs of days for each culture (excluding days with zero significant TE values). Specifically, in each scatter plot, the *x* value of a given point is the average inward TE from the associated node on an earlier day and the *y* value of that same point is the total outgoing TE from the same node but on a later day. The days in question are shown on the bottom and sides of the grids of scatter plots. The orange line shows the ordinary least squares regression. The Spearman correlation (*ρ*) between the outgoing TE values on the two days is displayed in each plot. Values of *ρ* significant at the 0.05 level are designated with an asterix and those significant at the 0.01 level are designated with a double asterix. A Bonferroni correction for multiple comparisons was used. (e) shows all recording day pairs for all cultures (where the pairs are always from the same culture) and the associated Spearman correlation between the outward TEs of nodes across this pair of recording days.

**TABLE VII:**
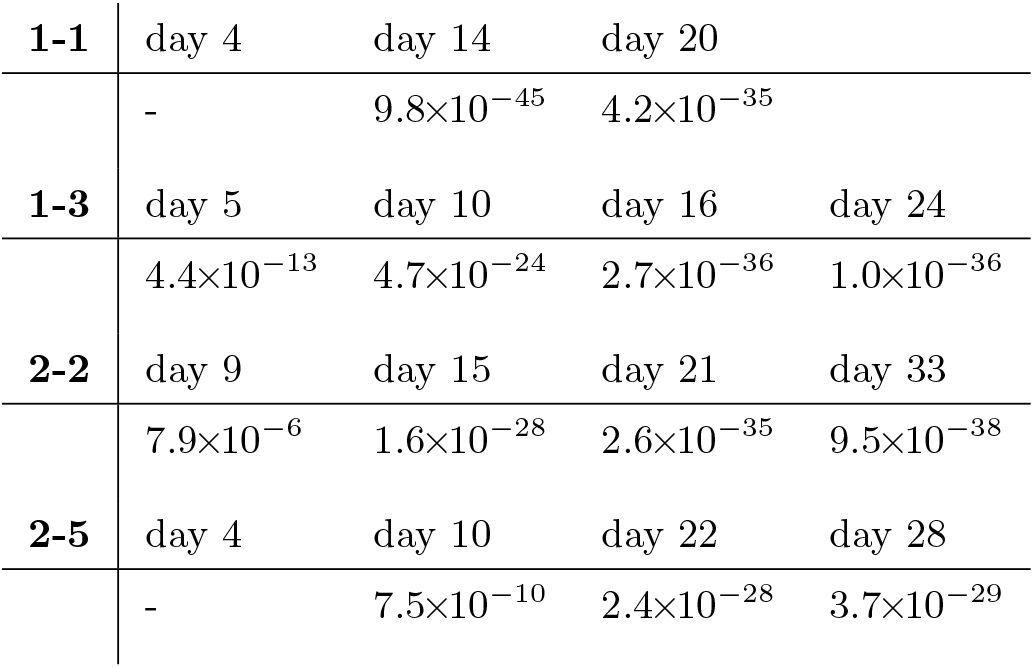
*p* values for the Shapiro-Wilke test [85] of normality for the distribution of TE values estimated in each recording. Only the statistically significant TE values are included in these tests. Recordings for which there were no statistically significant values estimated are left blank. These *p* values represent the probability that the associated test statistic is more extreme than that calculated on the estimated TE values, under the null hypothesis that these values are normally distributed. For any reasonable choice of *p* cutoff value, the null hypothesis is rejected in all recordings.

**TABLE VIII:**
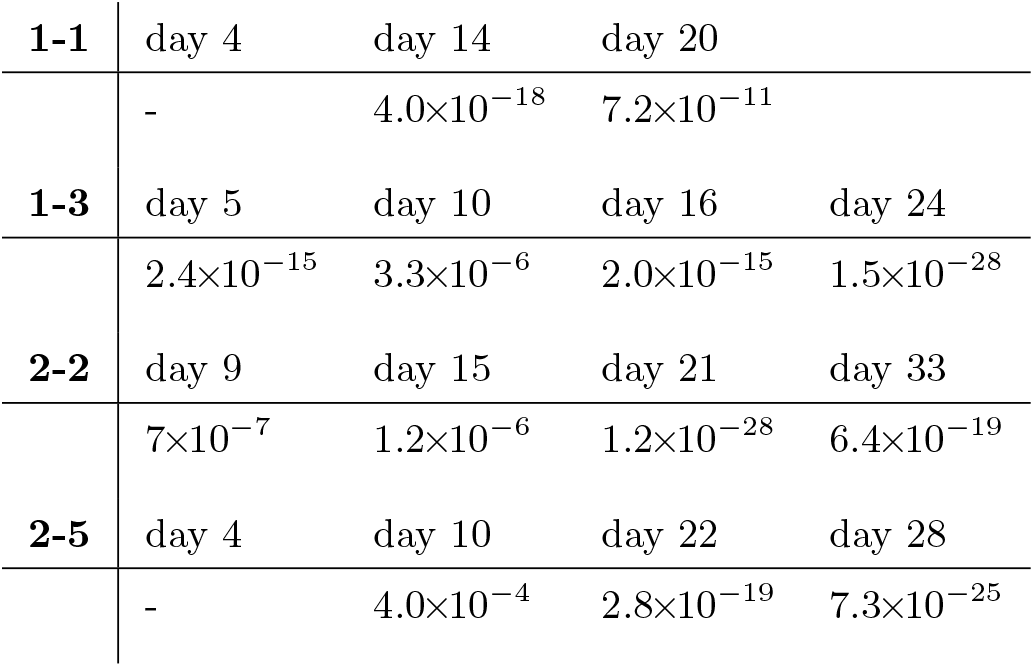
*p* values for the Shapiro-Wilke test [85] of log-normality for the distribution of TE values estimated in each recording. Only the statistically significant TE values are included in these tests. Recordings for which there were no statistically significant values estimated are left blank. These *p* values represent the probability that the associated test statistic is more extreme than that calculated on the logarithms of the estimated TE values, under the null hypothesis that these values are normally distributed. For any reasonable choice of *p* cutoff value, the null hypothesis is rejected in all recordings. It is interesting to note that the *p* values are often smaller on later days, despite the Q-Q plots in Fig. 9b suggesting the distribution is closer to log-normal. This is probably due to there being many more statistically significant TE values on these later days (see Table II).

Given that the distributions of the TE values were not well described by a log-normal distribution, we investigated the alternative that they could be described by a normal distribution. Fig. 9a displays Quantile-Quantile (QQ) plots [84] for the TE values against the normal distribution. In all recordings, the plotted points deviate substantially from the line *y* = *x*, indicating that the data is poorly described by a normal distribution. We also perform Shapiro-Wilke tests [85] for normality, the resulting *p* values are displayed in Table VII. The *p* values for every recording are incredibly low, meaning that we reject the null hypothesis of a normal distribution in every case.

These results contrast with observation of log-normal distributions of TE values in *in vitro* cultures of neurons [14, 15]. The difference may be due to the use of continuous-time estimator here in contrast to the discrete-time estimator used in previous studies. This estimator is more faithful to capturing the true underlying TE for spike trains (as per [22]), however it may be that the combination of the discrete-time estimator and use of only a single previous time-bin – in specifically *not* representing history dependence well – align more strongly with the component of the statistical relationship that follows a log-normal distribution. It is also possible that log-normal distributions of TE emerge later in development, and are simply not yet present in the early developmental stages observed here (noting that the fit to a log-normal distribution seems to improve for later DIV in Fig. 9b).

## Appendix B: Plots for Early Lock-in of Incoming TE

## References

[1] J. T. Lizier, M. Prokopenko, and A. Y. Zomaya, A framework for the local information dynamics of distributed computation in complex systems, in Guided self-organization: Inception accessed: (Springer, 2014) pp. 115–158.

[2] J. T. Lizier, The local information dynamics of distributed computation in complex systemsaccessed:, Springer Theses (Springer, Berlin / Heidelberg, 2013).

[3] T. Schreiber, Measuring information transfer, Physical review letters 85, 461 (2000).

[4] T. Bossomaier, L. Barnett, M. Harré, and J. T. Lizier, An introduction to transfer entropy, Cham: Springer International Publishing 65 (2016).

[5] J. G. Orlandi, O. Stetter, J. Soriano, T. Geisel, and D. Battaglia, Transfer entropy reconstruction and labeling of neuronal connections from simulated calcium imaging, PloS one 9, e98842 (2014).

[6] V. Maki-Marttunen, I. Diez, J. M. Cortes, D. R. Chialvo, and M. Villarreal, Disruption of transfer entropy and inter-hemispheric brain functional connectivity in patients with disorder of consciousness, Frontiers in neuroinformatics 7, 24 (2013).

[7] J. T. Lizier, J. Heinzle, A. Horstmann, J.-D. Haynes, and M. Prokopenko, Multivariate information-theoretic measures reveal directed information structure and task relevant changes in fmri connectivity, Journal of computational neuroscience 30, 85 (2011).

[8] M. Wibral, B. Rahm, M. Rieder, M. Lindner, R. Vicente, and J. Kaiser, Transfer entropy in magnetoencephalographic data: quantifying information flow in cortical and cerebellar networks, Progress in biophysics and molecular biology 105, 80 (2011).

[9] M. H. I. Shovon, N. Nandagopal, R. Vijayalakshmi, J. T. Du, and B. Cocks, Directed connectivity analysis of functional brain networks during cognitive activity using transfer entropy, Neural Processing Letters 45, 807 (2017).

[10] C.-S. Huang, N. R. Pal, C.-H. Chuang, and C.-T. Lin, Identifying changes in eeg information transfer during drowsy driving by transfer entropy, Frontiers in human neuroscience 9, 570 (2015).

[11] S. Stramaglia, G.-R. Wu, M. Pellicoro, and D. Marinazzo, Expanding the transfer entropy to identify information circuits in complex systems, Physical Review E 86, 066211 (2012).

[12] D. Marinazzo, O. Gosseries, M. Boly, D. Ledoux, M. Rosanova, M. Massimini, Q. Noirhomme, and S. Laureys, Directed information transfer in scalp electroencephalographic recordings: insights on disorders of consciousness, Clinical EEG and neuroscience 45, 33 (2014).

[13] M. Wibral, J. Lizier, S. Vogler, V. Priesemann, and R. Galuske, Local active information storage as a tool to understand distributed neural information processing, Frontiers in neuroinformatics 8, 1 (2014).

[14] S. Nigam, M. Shimono, S. Ito, F.-C. Yeh, N. Timme, M. Myroshnychenko, C. C. Lapish, Z. Tosi, P. Hottowy, W. C. Smith, et al., Rich-club organization in effective connectivity among cortical neurons, Journal of Neuroscience 36, 670 (2016).

[15] M. Shimono and J. M. Beggs, Functional clusters, hubs, and communities in the cortical microconnectome, Cerebral Cortex 25, 3743 (2015).

[16] E. Matsuda, T. Mita, J. Hubert, M. Oka, D. Bakkum, U. Frey, H. Takahashi, and T. Ikegami, Multiple time scales observed in spontaneously evolved neurons on high-density cmos electrode array, in Artificial Life Conference Proceedings 13 accessed: (MIT Press, 2013) pp. 1075–1082.

[17] N. Timme, S. Ito, M. Myroshnychenko, F.-C. Yeh, E. Hiolski, P. Hottowy, and J. M. Beggs, Multiplex networks of cortical and hippocampal neurons revealed at different timescales, PloS One 9, e115764 (2014).

[18] M. Kajiwara, R. Nomura, F. Goetze, M. Kawabata, Y. Isomura, T. Akutsu, and M. Shimono, Inhibitory neurons exhibit high controlling ability in the cortical microconnectome, PLOS Computational Biology 17, e1008846 (2021).

[19] N. M. Timme, S. Ito, M. Myroshnychenko, S. Nigam, M. Shimono, F.-C. Yeh, P. Hottowy, A. M. Litke, and J. M. Beggs, High-degree neurons feed cortical computations, PLoS Computational Biology 12, e1004858 (2016).

[20] M. Wibral, C. Finn, P. Wollstadt, J. T. Lizier, and V. Priesemann, Quantifying information modification in developing neural networks via partial information decomposition, Entropy 19, 494 (2017).

[21] D. A. Wagenaar, J. Pine, and S. M. Potter, An extremely rich repertoire of bursting patterns during the development of cortical cultures, BMC Neuroscience 7, 1 (2006).

[22] D. P. Shorten, R. E. Spinney, and J. T. Lizier, Estimating transfer entropy in continuous time between neural spike trains or other event-based data, PLOS Computational Biology 17, e1008054 (2021).

[23] M. S. Schroeter, P. Charlesworth, M. G. Kitzbichler, O. Paulsen, and E. T. Bullmore, Emergence of rich-club topology and coordinated dynamics in development of hippocampal functional networks in vitro, Journal of Neuroscience 35, 5459 (2015).

[24] Network activity of developing cortical cultures in vitro accessed:, http://neurodatasharing.bme.gatech.edu/development-data/html/index.html, accessed: 2021-01-03.

[25] V. Pasquale, P. Massobrio, L. Bologna, M. Chiappalone, and S. Martinoia, Self-organization and neuronal avalanches in networks of dissociated cortical neurons, Neuroscience 153, 1354 (2008).

[26] V. Priesemann, M. Wibral, M. Valderrama, R. Pröpper, M. Le Van Quyen, T. Geisel, J. Triesch, D. Nikolić, and M. H. Munk, Spike avalanches in vivo suggest a driven, slightly subcritical brain state, Frontiers in systems neuroscience 8, 108 (2014).

[27] V. Priesemann, M. Valderrama, M. Wibral, and M. Le Van Quyen, Neuronal avalanches differ from wakefulness to deep sleep–evidence from intracranial depth recordings in humans, PLoS Comput Biol 9, e1002985 (2013).

[28] V. Priesemann, M. H. Munk, and M. Wibral, Subsampling effects in neuronal avalanche distributions recorded in vivo, BMC neuroscience 10, 1 (2009).

[29] J. E. Lisman, Bursts as a unit of neural information: making unreliable synapses reliable, Trends in neurosciences 20, 38 (1997).

[30] R. Krahe and F. Gabbiani, Burst firing in sensory systems, Nature Reviews Neuroscience 5, 13 (2004).

[31] W. L. Shew, H. Yang, S. Yu, R. Roy, and D. Plenz, Information capacity and transmission are maximized in balanced cortical networks with neuronal avalanches, Journal of neuroscience 31, 55 (2011).

[32] O. Kinouchi and M. Copelli, Optimal dynamical range of excitable networks at criticality, Nature physics 2, 348 (2006).

[33] C. Haldeman and J. M. Beggs, Critical branching captures activity in living neural networks and maximizes the number of metastable states, Physical review letters 94, 058101 (2005).

[34] M. Rubinov, O. Sporns, J.-P. Thivierge, and M. Breakspear, Neurobiologically realistic determinants of self-organized criticality in networks of spiking neurons, PLoS Comput Biol 7, e1002038 (2011).

[35] B. Cramer, D. Stöckel, M. Kreft, M. Wibral, J. Schemmel, K. Meier, and V. Priesemann, Control of criticality and computation in spiking neuromorphic networks with plasticity, Nature communications 11, 1 (2020).

[36] E. Maeda, H. Robinson, and A. Kawana, The mechanisms of generation and propagation of synchronized bursting in developing networks of cortical neurons, Journal of Neuroscience 15, 6834 (1995).

[37] J. T. Lizier, M. Prokopenko, and A. Y. Zomaya, Local information transfer as a spatiotemporal filter for complex systems, Physical Review E 77, 026110 (2008).

[38] J. T. Lizier, Measuring the dynamics of information processing on a local scale in time and space, in Directed Information Measures in Neuroscience, Understanding Complex Systems, edited by M. Wibral, R. Vicente, and J. T. Lizier (Springer, Berlin/Heidelberg, 2014) pp. 161–193.

[39] M. Wibral, R. Vicente, and J. T. Lizier, Directed information measures in neuroscience accessed: (Springer, 2014).

[40] M. Khoshkhou and A. Montakhab, Spike-timing-dependent plasticity with axonal delay tunes networks of izhikevich neurons to the edge of synchronization transition with scale-free avalanches, Frontiers in systems neuroscience 13, 73 (2019).

[41] E. M. Izhikevich, Simple model of spiking neurons, IEEE Transactions on neural networks 14, 1569 (2003).

[42] N. Caporale and Y. Dan, Spike timing–dependent plasticity: a hebbian learning rule, Annu. Rev. Neurosci. 31, 25 (2008).

[43] R. Zeraati, V. Priesemann, and A. Levina, Self-organization toward criticality by synaptic plasticity, Frontiers in Physics 9, 103 (2021).

[44] J. T. Lizier, M. Prokopenko, and A. Y. Zomaya, Information modification and particle collisions in distributed computation, Chaos: An Interdisciplinary Journal of Nonlinear Science 20, 037109 (2010).

[45] J. T. Lizier, M. Prokopenko, and A. Y. Zomaya, Local measures of information storage in complex distributed computation, Information Sciences 208, 39 (2012).

[46] M. Li, Y. Han, M. J. Aburn, M. Breakspear, R. A. Poldrack, J. M. Shine, and J. T. Lizier, Transitions in information processing dynamics at the whole-brain network level are driven by alterations in neural gain, PLoS computational biology 15, e1006957 (2019).

[47] D. Marinazzo, G. Wu, M. Pellicoro, L. Angelini, and S. Stramaglia, Information flow in networks and the law of diminishing marginal returns: evidence from modeling and human electroencephalographic recordings, PLoS one 7, e45026 (2012).

[48] D. Marinazzo, M. Pellicoro, G. Wu, L. Angelini, J. M. Cortés, and S. Stramaglia, Information transfer and criticality in the ising model on the human connectome, PloS one 9, e93616 (2014).

[49] R. V. Ceguerra, J. T. Lizier, and A. Y. Zomaya, Information storage and transfer in the synchronization process in locally-connected networks, in 2011 IEEE Symposium on Artificial Life (ALIFE) (IEEE, 2011) pp. 54–61.

[50] L. Novelli, F. M. Atay, J. Jost, and J. T. Lizier, Deriving pairwise transfer entropy from network structure and motifs, Proceedings of the Royal Society A: Mathematical, Physical and Engineering Sciences 476, 20190779 (2020).

[51] R. H. Goodman and M. Porfiri, Topological features determining the error in the inference of networks using transfer entropy, Mathematics in Engineering 2, 34 (2019).

[52] J. T. Lizier, M. Prokopenko, and D. J. Cornforth, The information dynamics of cascading failures in energy networks, in Proceedings of the European Conference on Complex Systems (ECCS), Warwick, UK accessed: (Citeseer, 2009) p. 54.

[53] M. Gilson, A. N. Burkitt, D. B. Grayden, D. A. Thomas, and J. L. van Hemmen, Emergence of network structure due to spike-timing-dependent plasticity in recurrent neuronal networks. ii. input selectivitysymmetry breaking, Biological Cybernetics 101, 103 (2009).

[54] S. Kunkel, M. Diesmann, and A. Morrison, Limits to the development of feed-forward structures in large recurrent neuronal networks, Frontiers in computational neuroscience 4, 160 (2011).

[55] M. A. Frost and R. Goebel, Measuring structural–functional correspondence: spatial variability of specialised brain regions after macro-anatomical alignment, Neuroimage 59, 1369 (2012).

[56] J. R. Cohen and M. D’Esposito, The segregation and integration of distinct brain networks and their relationship to cognition, Journal of Neuroscience 36, 12083 (2016).

[57] D. A. Wagenaar, Z. Nadasdy, and S. M. Potter, Persistent dynamic attractors in activity patterns of cultured neuronal networks, Physical Review E 73, 051907 (2006).

[58] D. S. Bassett and O. Sporns, Network neuroscience, Nature neuroscience 20, 353 (2017).

[59] L. Novelli and J. T. Lizier, Inferring network properties from time series using transfer entropy and mutual information: Validation of multivariate versus bivariate approaches, Network Neuroscience 5, 373 (2021).

[60] L. Novelli, P. Wollstadt, P. Mediano, M. Wibral, and J. T. Lizier, Large-scale directed network inference with multivariate transfer entropy and hierarchical statistical testing, Network Neuroscience 3, 827 (2019).

[61] A. Kraskov, H. Stögbauer, and P. Grassberger, Estimating mutual information, Physical review E 69, 066138 (2004).

[62] D. J. MacKay and D. J. Mac Kay, Information theory, inference and learning algorithms accessed: (Cambridge university press, 2003).

[63] R. E. Spinney and J. T. Lizier, Characterizing information-theoretic storage and transfer in continuous time processes, Physical Review E 98, 012314 (2018).

[64] G. Mijatovic, Y. Antonacci, T. L. Turukalo, L. Minati, and L. Faes, An information-theoretic framework to measure the dynamic interaction between neural spike trains, IEEE Transactions on Biomedical Engineering (2021).

[65] L. Kozachenko and N. N. Leonenko, Sample estimate of the entropy of a random vector, Problemy Peredachi Informatsii 23, 9 (1987).

[66] M. Garofalo, T. Nieus, P. Massobrio, and S. Martinoia, Evaluation of the performance of information theory-based methods and cross-correlation to estimate the functional connectivity in cortical networks, PloS One 4, e6482 (2009).

[67] M. Kajiwara, R. Nomura, F. Goetze, T. Akutsu, and M. Shimono, Inhibitory neurons are a central controlling regulator in the effective cortical microconnectome., bioRxiv (2020).

[68] J. W. Aldridge and S. Gilman, The temporal structure of spike trains in the primate basal ganglia: afferent regulation of bursting demonstrated with precentral cerebral cortical ablation, Brain Research 543, 123 (1991).

[69] L. Rudelt, D. G. Marx, M. Wibral, and V. Priesemann, Embedding optimization reveals long-lasting history dependence in neural spiking activity, PLOS Computational Biology 17, e1008927 (2021).

[70] I. Nemenman, G. D. Lewen, W. Bialek, and R. R. D. R. Van Steveninck, Neural coding of natural stimuli: information at sub-millisecond resolution, PLoS Computational Biology 4, e1000025 (2008).

[71] C. Kayser, N. K. Logothetis, and S. Panzeri, Millisecond encoding precision of auditory cortex neurons, Proceedings of the National Academy of Sciences 107, 16976 (2010).

[72] S. J. Sober, S. Sponberg, I. Nemenman, and L. H. Ting, Millisecond spike timing codes for motor control, Trends in Neurosciences 41, 644 (2018).

[73] J. A. Garcia-Lazaro, L. A. Belliveau, and N. A. Lesica, Independent population coding of speech with sub-millisecond precision, Journal of Neuroscience 33, 19362 (2013).

[74] E. Y. Erten, J. T. Lizier, M. Piraveenan, and M. Prokopenko, Criticality and information dynamics in epidemiological models, Entropy 19, 194 (2017).

[75] J. Garland, R. G. James, and E. Bradley, Leveraging information storage to select forecast-optimal parameters for delay-coordinate reconstructions, Physical Review E 93, 022221 (2016).

[76] M. Hansen, A. Burns, C. Monk, C. Schutz, J. Lizier, I. Ramnarine, A. Ward, and J. Krause, The effect of predation risk on group behaviour and information flow during repeated collective decisions, Animal Behaviour 173, 215 (2021).

[77] P. Wollstadt, J. T. Lizier, R. Vicente, C. Finn, M. Martinez-Zarzuela, P. Mediano, L. Novelli, and M. Wibral, Idtxl: The information dynamics toolkit xl: a python package for the efficient analysis of multivariate information dynamics in networks, arXiv preprint arXiv:1807.10459 (2018).

[78] A. Zalesky, A. Fornito, L. Cocchi, L. L. Gollo, M. P. van den Heuvel, and M. Breakspear, Connectome sensitivity or specificity: which is more important?, Neuroimage 142, 407 (2016).

[79] G. Rupert Jr et al., Simultaneous statistical inference accessed: (Springer Science & Business Media, 2012).

[80] Y. Kaneoke and J. Vitek, Burst and oscillation as disparate neuronal properties, Journal of neuroscience methods 68, 211 (1996).

[81] D. Wagenaar, T. B. DeMarse, and S. M. Potter, Meabench: A toolset for multi-electrode data acquisition and on-line analysis, in Conference Proceedings. 2nd International IEEE EMBS Conference on Neural Engineering, 2005. accessed: (IEEE, 2005) pp. 518–521.

[82] J. V. Selinger, N. V. Kulagina, T. J. O’Shaughnessy, W. Ma, and J. J. Pancrazio, Methods for characterizing interspike intervals and identifying bursts in neuronal activity, Journal of neuroscience methods 162, 64 (2007).

[83] D. J. Bakkum, M. Radivojevic, U. Frey, F. Franke, A. Hierlemann, and H. Takahashi, Parameters for burst detection, Frontiers in computational neuroscience 7, 193 (2014).

[84] J. D. Gibbons and S. Chakraborti, Nonparametric statistical inference accessed: (CRC press, 2020).

[85] S. S. Shapiro and M. B. Wilk, An analysis of variance test for normality (complete samples), Biometrika 52, 591 (1965).

